# *Salmonella* effectors SseK1 and SseK3 target death domain proteins in the TNF and TRAIL signaling pathways

**DOI:** 10.1101/359117

**Authors:** Joshua P M Newson, Nichollas E Scott, Ivy Yeuk Wah Chung, Tania Wong Fok Lung, Cristina Giogha, Nancy Wang, Richard A Strugnell, Nat F Brown, Miroslaw Cygler, Jaclyn S Pearson, Elizabeth L Hartland

## Abstract

Strains of *Salmonella* utilise two distinct type three secretion systems to deliver effector proteins directly into host cells. The *Salmonella* effectors SseK1 and SseK3 are arginine glycosyltransferases that modify mammalian death domain containing proteins with N-acetyl glucosamine (GlcNAc) when overexpressed ectopically or as recombinant protein fusions. Here, we combined Arg-GlcNAc glycopeptide immunoprecipitation and mass spectrometry to identify host proteins GlcNAcylated by endogenous levels of SseK1 and SseK3 during *Salmonella* infection. We observed that SseK1 modified the mammalian signaling protein TRADD, but not FADD as previously reported. Overexpression of SseK1 greatly broadened substrate specificity, while ectopic co-expression of SseK1 and TRADD increased the range of modified arginine residues within the death domain of TRADD. In contrast, endogenous levels of SseK3 resulted in modification of the death domains of receptors of the mammalian TNF superfamily, TNFR1 and TRAILR, at residues Arg^376^ and Arg^293^ respectively. Structural studies on SseK3 showed that the enzyme displays a classic GT-A glycosyltransferase fold and binds UDP-GlcNAc in a narrow and deep cleft with the GlcNAc facing the surface. Together our data suggests that Salmonellae carrying *sseK1* and *sseK3* employ the glycosyltransferase effectors to antagonise different components of death receptor signaling.

## Author Summary

Many Gram-negative pathogens employ type three secretion systems to translocate specialised bacterial effector proteins into the host cell. The activities of many *Salmonella* effectors are not well understood, and identifying the host targets of these effectors will lead to a better understanding of *Salmonella* pathogenesis. The overexpression of effectors *in vitro* can provide useful insights into their function, but these results may not represent the true biological role of these effectors during infection. In this study, we showed that endogenous levels of two *Salmonella* effectors target different host death domain containing signaling proteins that are required for inflammatory and cell death signaling. The study developed new tools to study cognate effector interactions and established the important of effector expression levels in defining natural and unnatural interactions.

## Introduction

Pathogenic serovars of *Salmonella* utilise two type three secretion systems (T3SS), encoded by *Salmonella* pathogenicity island-1 and −2 (SPI-1 and SPI-2) to deliver distinct cohorts of effector proteins into host cells during infection [1, 2]. These effector proteins subvert normal cellular processes and collectively enable the bacteria to invade and persist within host cells, partially through the manipulation of inflammatory cell signaling and programmed cell death (reviewed in [3, 4]). While the importance of effector translocation to pathogenesis is well established, the specific contribution of many effectors is still unclear. In particular, many effectors translocated by the SPI-2 encoded T3SS remain poorly characterised.

SseK1, SseK2, and SseK3 comprise a family of highly similar *Salmonella* effectors that are translocated by the SPI-2 T3SS during infection [5, 6]. SseK family members show high sequence similarity to NleB1, a T3SS effector protein from enteropathogenic *Escherichia coli* (EPEC), which functions as an arginine glycosyltransferase and catalyses the addition of *N*-acetylglucosamine (GlcNAc) to arginine residues of the mammalian signaling adaptors FADD and TRADD, a modification termed Arg-GlcNAcylation [7, 8]. A recent report provides evidence that SseK1 and SseK3, but not SseK2, also function as Arg-GlcNAc glycosyltransferases [9]. Mutation of a conserved DxD catalytic motif within SseK1 and SseK3 abrogates their glycosyltransferase activity [9], consistent with findings for NleB1 [7, 8]. These studies describing catalytically important regions of the glycosyltransferases provide opportunities to better understand the function of these novel enzymes [10].

Although well recognised as glycosyltransferases, there are conflicting reports regarding the host substrates of the SseK effectors. One report suggested that recombinant SseK1 modifies recombinant TRADD *in vitro* [8], whereas a subsequent report suggested SseK1 modifies both TRADD and FADD when co-expressed ectopically in mammalian cell lines [9]. Yet another report using *in vitro* glycosylation assays suggested that recombinant SseK1 glycosylates GAPDH but not FADD [11]. SseK3 on the other hand was reported to bind but not modify the E3-ubiquitin ligase TRIM32 [12], and was shown to weakly modify TRADD but not FADD [9]. While these studies provide useful insights, the true role of these effectors is better interrogated through non-biased screens conducted under conditions that reflect endogenous levels of the effectors and host proteins [13].

Here, we explored the endogenous Arg-GlcNAc glycosyltransferase activity of SseK1 and SseK3 during *Salmonella* infection. Using a mass spectrometry-based approach to enrich arginine GlcNAcylated peptides from infected host cells [13], we found that SseK1 modified the signaling adaptor TRADD, while SseK3 modified the signaling receptors TNFR1 and TRAILR. In addition, we performed structural studies on SseK3 and showed that the enzyme displays a classic GT-A glycosyltransferase fold and binds UDP-GlcNAc in a narrow and deep cleft with the GlcNAc facing the surface. Together these studies suggest that *Salmonella* has evolved multiple means to manipulate death receptor signaling through the acquisition of arginine glycosyltransferases with differing substrate specificities.

## Results

### Mutation of a conserved glutamic acid abrogates the catalytic activity of SseK1 and SseK3

Previous reports have indicated that SseK1 and SseK3 function as arginine glycosyltransferases which modify a conserved arginine residue in the death domains of several mammalian immune signaling proteins [8, 9]. Mutation of a conserved DxD catalytic motif within SseK1 (Asp^223^-Ala^224-^Asp^225^) and SseK3 (Asp^226^-Ala^227^-Asp^228^) abrogated their glycosyltransferase activity [9], consistent with findings for the homologous T3SS effector, NleB1, from EPEC [7, 8]. Mutation of a single glutamic acid (Glu^253^) also impairs the ability of NleB1 to inhibit NF-κB activation following TNF stimulation of mammalian cells [8], and renders NleB1 catalytically inactive [10]. Here, we confirmed the importance of this conserved glutamic acid to the biochemical activity of SseK1 and SseK3 by infecting RAW264.7 cells with a *Salmonella* Typhimurium SL1344 triple *ΔsseK123* deletion mutant [6] complemented with either native SseK1, −2 or −3, catalytic triad mutants, or mutants lacking the conserved glutamic acid (SseK1_E255A_, SseK2_E271A_, and SseK3_E258A_). Cell lysates were assessed by immunoblot using an antibody specific for arginine glycosylation (Fig. 1A and B). Native SseK1 and SseK3, but not SseK2, catalysed Arg-GlcNAcylation, while all catalytic triad mutants and glutamic acid mutants showed no activity. Notably, overexpression of both SseK1 and SseK3 increased levels of arginine glycosylation relative to wild-type SL1344, suggesting that overexpression may cause non-authentic Arg-GlcNAcylation of host substrates.

**Figure 1.**
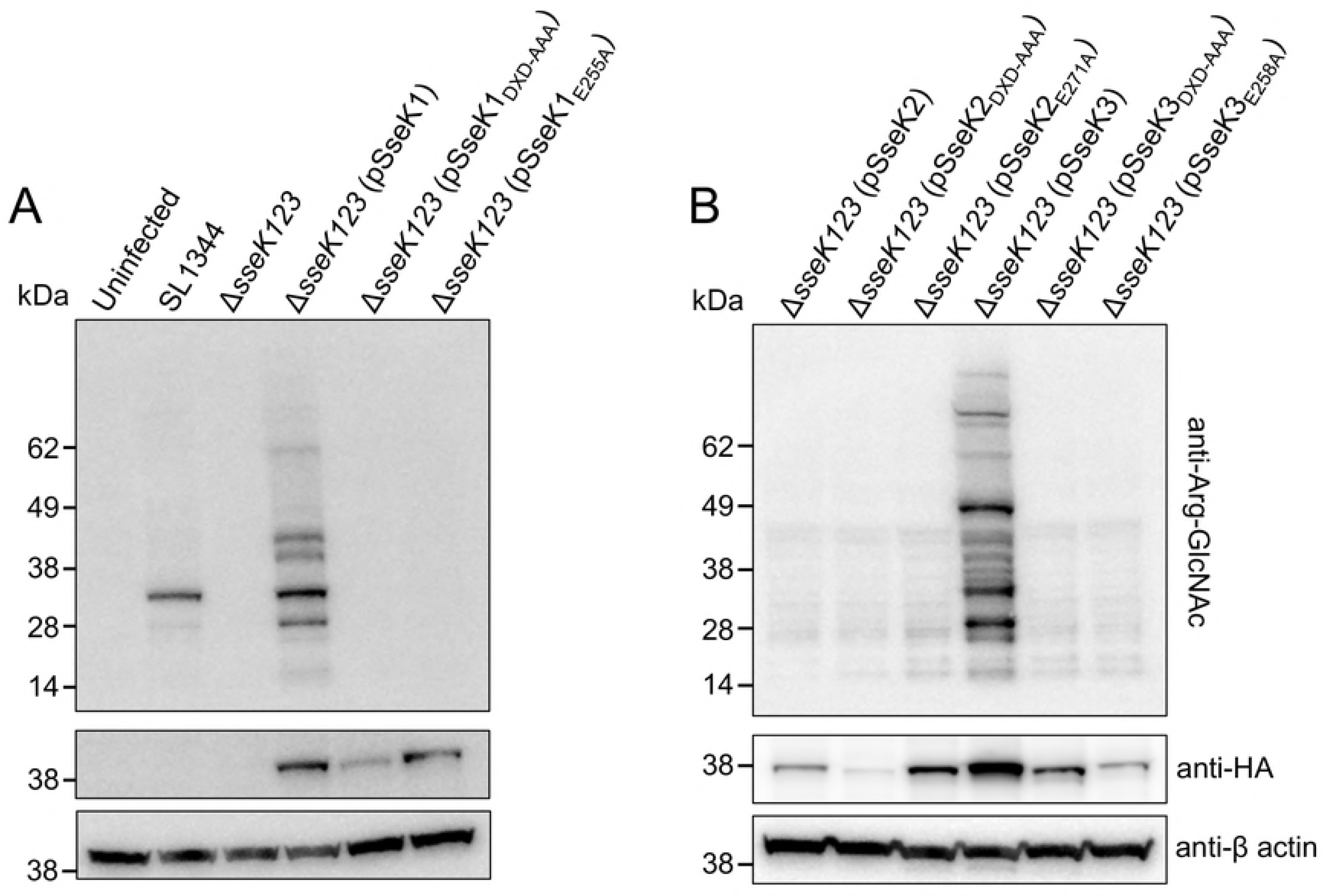
**Immunoblot of RAW264.7 cells infected with derivatives of** ***S.*** **Typhimurium SL1344.** Wild type *S.* Typhimurium, a triple Δ*ssek123* mutant and triple mutant complemented with plasmids encoding one of HA-tagged SseK1, SseK2 or SseK3, or with catalytically-inactive effector derivatives were used to infect RAW264.7 for 20 h, as indicated (A and B). Overexpression of the effectors was induced during host cell infection by the addition of 1 mM IPTG. RAW264.7 cells were lysed and proteins detected by immunoblot with anti-ArgGlcNAc, anti-HA, and anti-β-actin antibodies as indicated. Representative immunoblot of at least three independent experiments.

### The host signaling adaptor TRADD is the preferred substrate of SseK1

A number of reports have used *in vitro* experiments to identify possible substrates of SseK1 [8, 9, 11]. While these studies have demonstrated the ability of SseK1 to GlcNAcylate a given target, the approach is substrate specific and there requires knowledge of the potential targets. To identify the full range of host Arg-GlcNAcylated substrates, we developed a quantitative approach using mass spectrometry (MS) to enrich for arginine glycosylated peptides [13]. Using peptides derived from host cells infected with *S.* Typhimurium *ΔsseK123* over-expressing either SseK1 or inactive SseK1_E255A_, label-free MS based quantification revealed a broad range of substrates that were only modified in the presence of active SseK1 (Fig. 2A, Supplementary Table 1, Supplementary Fig. 1A). Among these substrates, mouse TRADD (mTRADD) was Arg-GlcNAcylated at Arg^243^ (Fig. 2B), which represented a novel site of modification. Unexpectedly, we also detected Arg-GlcNAcylation of a large number of *S.* Typhimurium proteins, among them the two-component response regulators OmpR and ArcA, as well as the ribosome-associated proteins RpsA and RbfA, and transcription-associated proteins RpoD and NusA.

**Figure 2.**
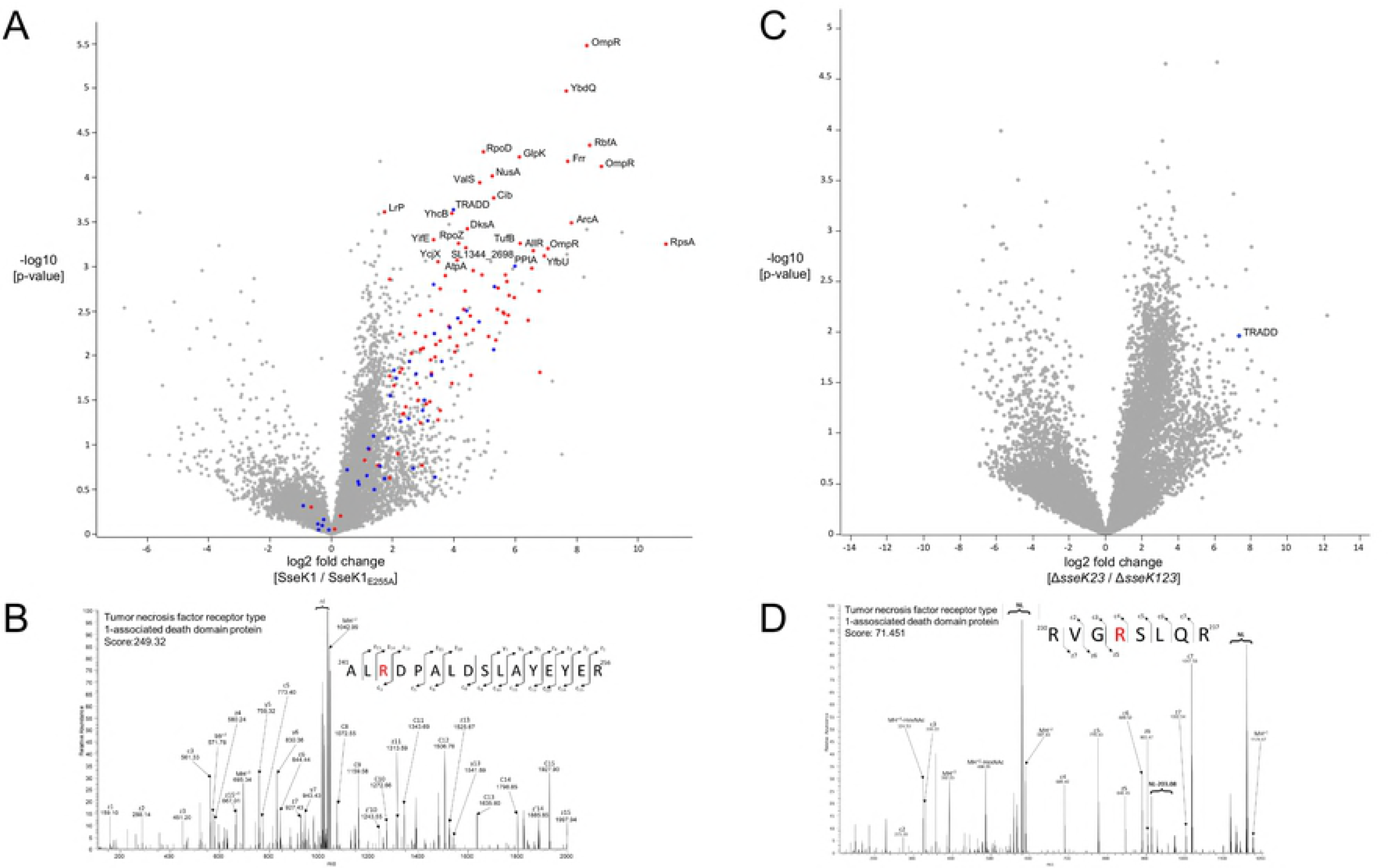
**Enrichment of peptides Arg-GlcNAcylated by SseK1 derived from** ***Salmonella-*****infected RAW264.7 cells.** (A) Label-free quantification of Arg-GlcNAc peptides immunoprecipitated from RAW264.7 cells infected with *S.* Typhimurium Δ*sseK123* complemented with either SseK1-HA or SseK1E_255A_-HA. Arg-GlcNAcylated peptides are presented as a volcano plot depicting mean ion intensity peptide ratios of SseK1-HA versus SseK1E_255A_-HA plotted against logarithmic *t* test *p* values from biological triplicate experiments. Arg-GlcNAcylated peptides with corresponding *t* test *p* values below 0.001 are annotated by protein name, with human peptides shaded blue and bacterial peptides shaded red. (B) Manually curated EThcD spectra showing glycosylation of Arg^243^ within the death domain of mouse TRADD, Andromeda score 249.32. Within MS/MS spectra NL denote neutral loss associated ions. (C) Parallel reaction monitoring of ArgGlcNAc peptide immunoprecipitated from RAW264.7 cells infected with *S.* Typhimurium Δ*sseK123* or *S.* Typhimurium Δ*sseK23*. Arginine-glycosylated peptides are presented as a volcano plot depicting mean log2 ion intensity peptide ratios of Δ*sseK123* versus Δ*sseK23* plotted against logarithmic *t* test *p* values from biological triplicate experiments. Arg-GlcNAcylated peptides are annotated by protein name and shaded blue. (D) Manually curated EThcD spectra showing glycosylation of Arg^233^ within the death domain of mouse TRADD, Andromeda score 71.45. Within MS/MS spectra NL denote neutral loss associated ions.

**Table 1.**
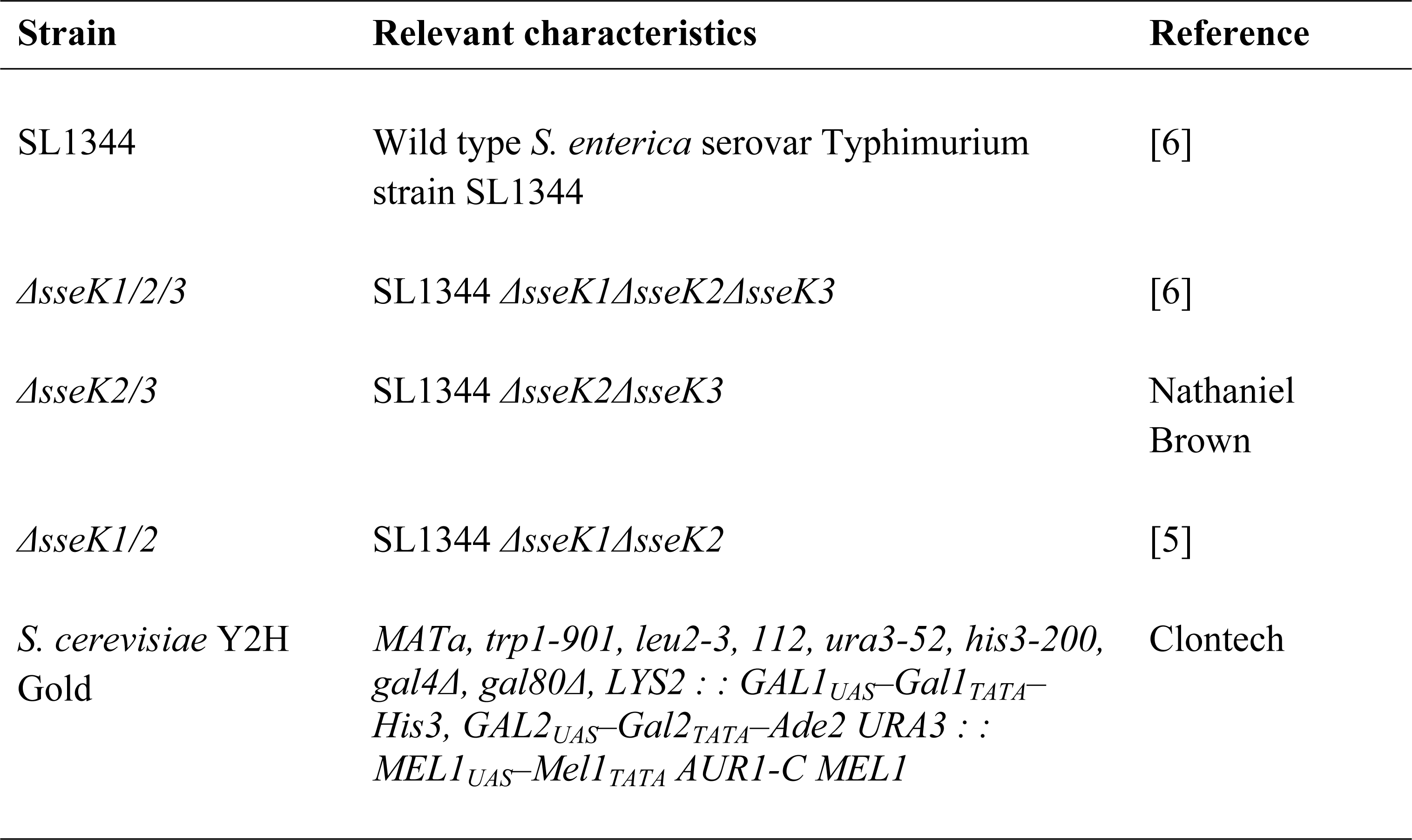
Strains used in this study.

We have previously shown that over-expression of NleB1 leads to enhanced levels of cellular arginine glycosylation [13]. Given that the overexpression of SseK1 and SseK3 also appeared to increase the range of substrates relative to endogenous wild type levels of expression (Fig 1A and B), we explored the activity of endogenous SseK1 during *S.* Typhimurium infection of RAW264.7 cells using our strategy for enrichment of Arg-GlcNAcylated peptides [13]. Here we compared Arg-GlcNAcylation during infection with a *ΔsseK23* double deletion mutant versus a *ΔsseK123* triple deletion mutant [6]. In addition to data dependent MS acquisition to enhance the detection of Arg-GlcNAcylation events, we applied parallel reaction monitoring, a targeted high-resolution MS approach [14], to specifically observe the Arg-GlcNAcylated species of murine TRADD (mTRADD) and FADD (mFADD) (Fig. 2C, Supplementary Table 2, Supplementary Fig. 1B). Using this approach, we found that mTRADD was glycosylated at Arg^233^ (Fig. 2D). Arg^233^ is the equivalent of the previously reported Arg^235^ of human TRADD (hTRADD) which is GlcNAcylated by NleB1 from EPEC *in vitro* [8]. There was no evidence of Arg-GlcNAcylated mFADD, despite the fact that mFADD was readily detectable within the input control samples (Supplementary Table 3, Supplementary Fig. 2). This suggested TRADD was the preferred substrate when SseK1 was translocated at native levels during infection.

**Table 2.**
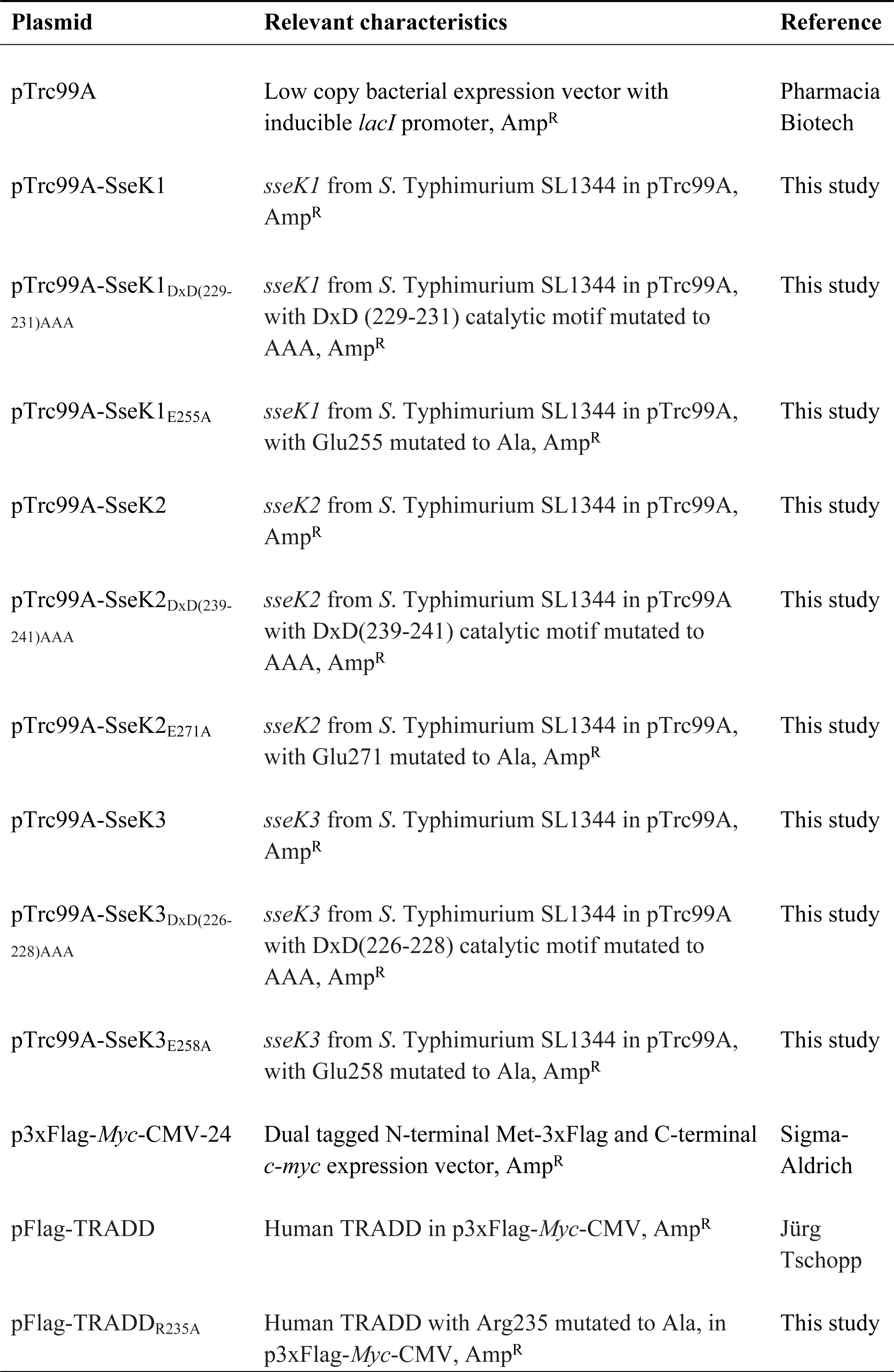
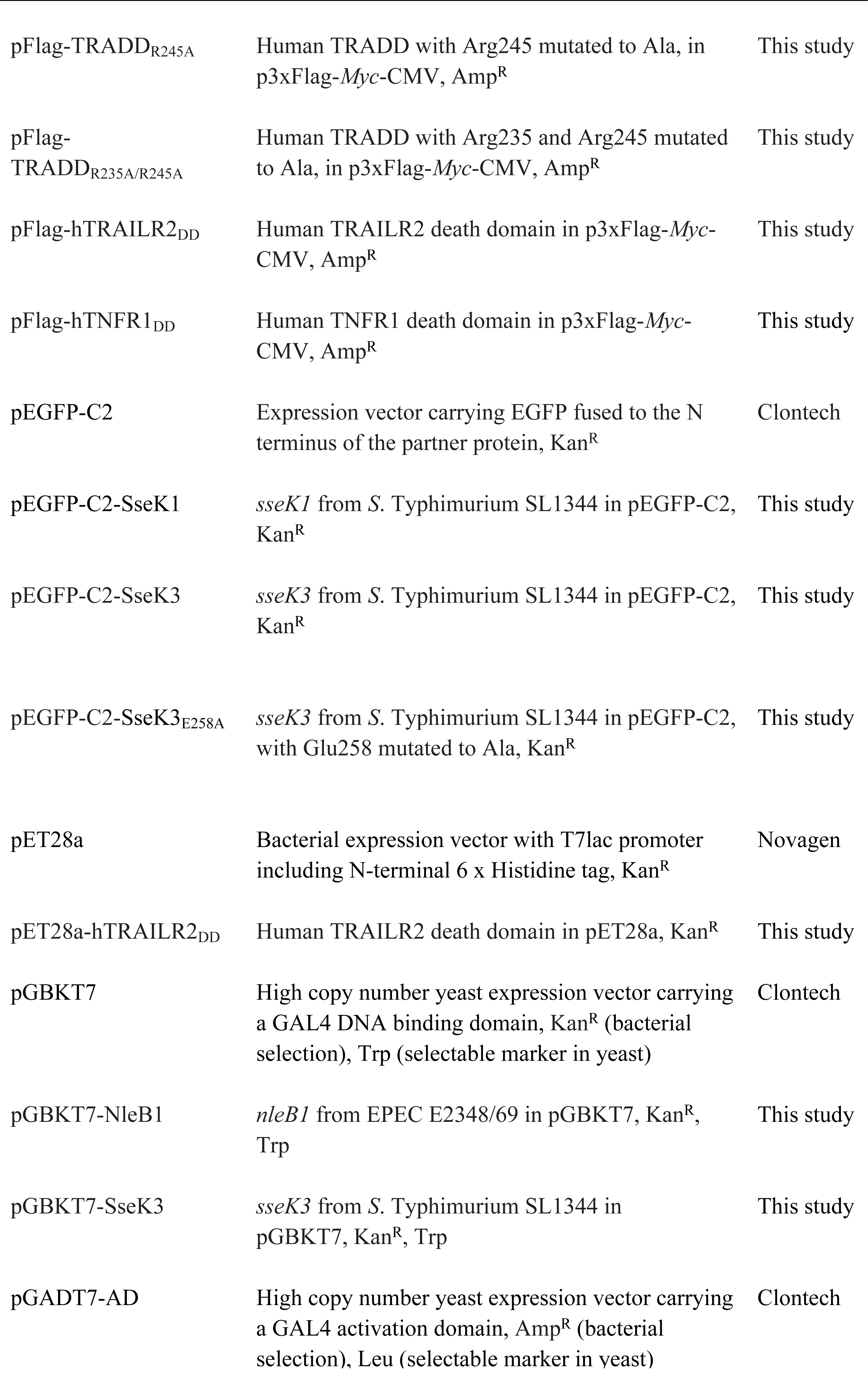
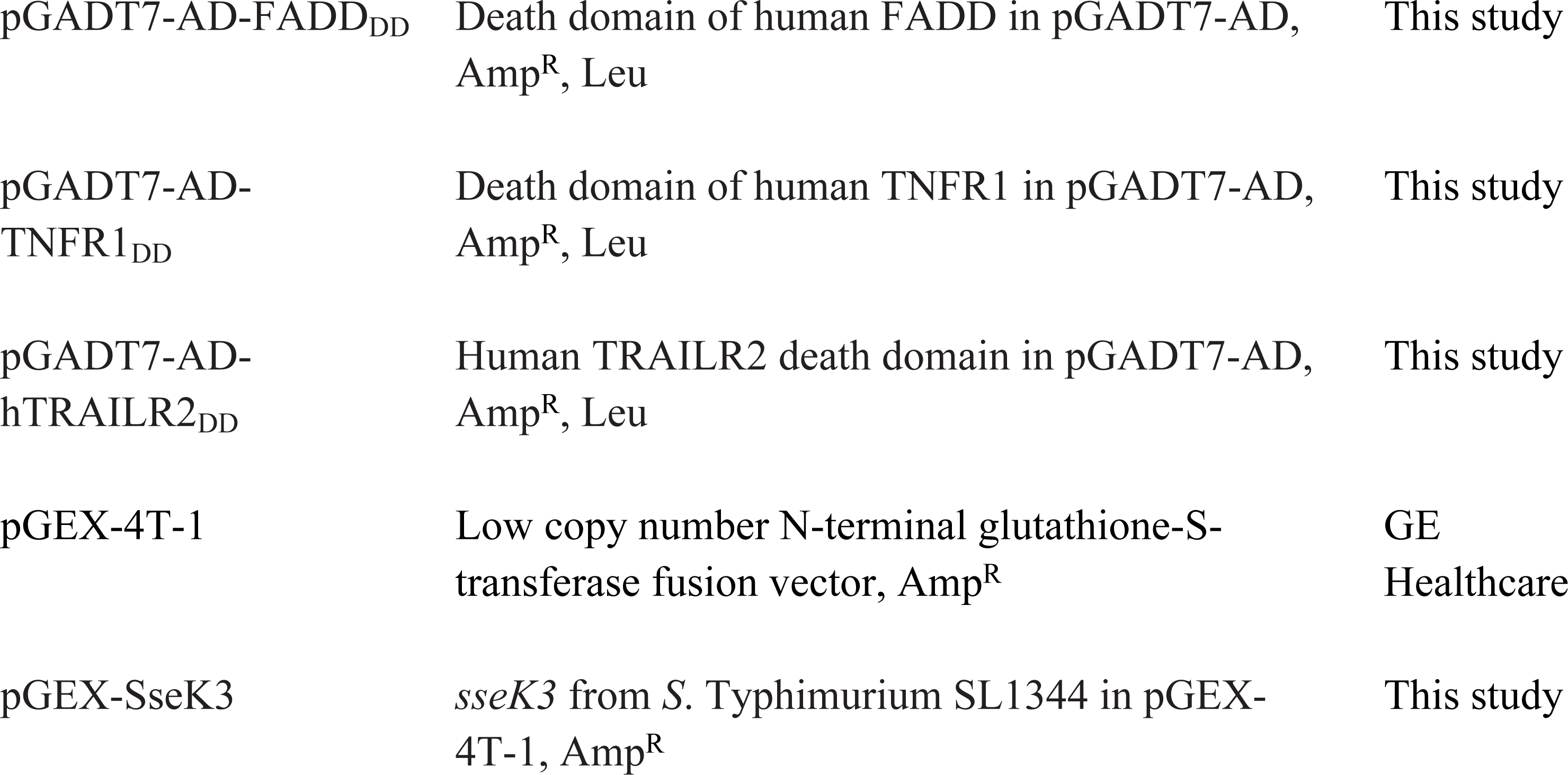
Plasmids used in this study.

| Plasmid | Relevant characteristics | Reference |
| --- | --- | --- |
| pTrc99A | Low copy bacterial expression vector with inducible *lacI* promoter, Amp<sup>R</sup> | Pharmacia Biotech |
| pTrc99A-SseK1 | *sseK1* from *S. Typhimurium* SL1344 in pTrc99A, Amp<sup>R</sup> | This study |
| pTrc99A-SseK1<sub>DxD(229-231)AAA</sub> | *sseK1* from *S. Typhimurium* SL1344 in pTrc99A, with DxD (229-231) catalytic motif mutated to AAA, Amp<sup>R</sup> | This study |
| pTrc99A-SseK1<sub>E255A</sub> | *sseK1* from *S. Typhimurium* SL1344 in pTrc99A, with Glu255 mutated to Ala, Amp<sup>R</sup> | This study |
| pTrc99A-SseK2 | *sseK2* from *S. Typhimurium* SL1344 in pTrc99A, Amp<sup>R</sup> | This study |
| pTrc99A-SseK2<sub>DxD(239-241)AAA</sub> | *sseK2* from *S. Typhimurium* SL1344 in pTrc99A with DxD(239-241) catalytic motif mutated to AAA, Amp<sup>R</sup> | This study |
| pTrc99A-SseK2<sub>E271A</sub> | *sseK2* from *S. Typhimurium* SL1344 in pTrc99A, with Glu271 mutated to Ala, Amp<sup>R</sup> | This study |
| pTrc99A-SseK3 | *sseK3* from *S. Typhimurium* SL1344 in pTrc99A, Amp<sup>R</sup> | This study |
| pTrc99A-SseK3<sub>DxD(226-228)AAA</sub> | *sseK3* from *S. Typhimurium* SL1344 in pTrc99A with DxD(226-228) catalytic motif mutated to AAA, Amp<sup>R</sup> | This study |
| pTrc99A-SseK3<sub>E258A</sub> | *sseK3* from *S. Typhimurium* SL1344 in pTrc99A, with Glu258 mutated to Ala, Amp<sup>R</sup> | This study |
| p3xFlag-*Myc*-CMV-24 | Dual tagged N-terminal Met-3xFlag and C-terminal *c-myc* expression vector, Amp<sup>R</sup> | Sigma-Aldrich |
| pFlag-TRADD | Human TRADD in p3xFlag-*Myc*-CMV, Amp<sup>R</sup> | Jürg Tschopp |
| pFlag-TRADD<sub>R235A</sub> | Human TRADD with Arg235 mutated to Ala, in p3xFlag-*Myc*-CMV, Amp<sup>R</sup> | This study |
| pFlag-TRADD<sub>R245A</sub> | Human TRADD with Arg245 mutated to Ala, in p3xFlag-*Myc*-CMV, Amp<sup>R</sup> | This study |
| pFlag-TRADD<sub>R235A/R245A</sub> | Human TRADD with Arg235 and Arg245 mutated to Ala, in p3xFlag-*Myc*-CMV, Amp<sup>R</sup> | This study |
| pFlag-hTRAILR2<sub>DD</sub> | Human TRAILR2 death domain in p3xFlag-*Myc*-CMV, Amp<sup>R</sup> | This study |
| pFlag-hTNFR1<sub>DD</sub> | Human TNFR1 death domain in p3xFlag-*Myc*-CMV, Amp<sup>R</sup> | This study |
| pEGFP-C2 | Expression vector carrying EGFP fused to the N terminus of the partner protein, Kan<sup>R</sup> | Clontech |
| pEGFP-C2-SseK1 | *sseK1* from *S. Typhimurium* SL1344 in pEGFP-C2, Kan<sup>R</sup> | This study |
| pEGFP-C2-SseK3 | *sseK3* from *S. Typhimurium* SL1344 in pEGFP-C2, Kan<sup>R</sup> | This study |
| pEGFP-C2-SseK3<sub>E258A</sub> | *sseK3* from *S. Typhimurium* SL1344 in pEGFP-C2, with Glu258 mutated to Ala, Kan<sup>R</sup> | This study |
| pET28a | Bacterial expression vector with T7lac promoter including N-terminal 6 x Histidine tag, Kan<sup>R</sup> | Novagen |
| pET28a-hTRAILR2<sub>DD</sub> | Human TRAILR2 death domain in pET28a, Kan<sup>R</sup> | This study |
| pGBKT7 | High copy number yeast expression vector carrying a GAL4 DNA binding domain, Kan<sup>R</sup> (bacterial selection), Trp (selectable marker in yeast) | Clontech |
| pGBKT7-NleB1 | *nleB1* from EPEC E2348/69 in pGBKT7, Kan<sup>R</sup>, Trp | This study |
| pGBKT7-SseK3 | *sseK3* from *S. Typhimurium* SL1344 in pGBKT7, Kan<sup>R</sup>, Trp | This study |
| pGADT7-AD | High copy number yeast expression vector carrying a GAL4 activation domain, Amp<sup>R</sup> (bacterial selection), Leu (selectable marker in yeast) | Clontech |
| pGADT7-AD-FADD<sub>DD</sub> | Death domain of human FADD in pGADT7-AD, Amp<sup>R</sup>, Leu | This study |
| pGADT7-AD-TNFR1<sub>DD</sub> | Death domain of human TNFR1 in pGADT7-AD, Amp<sup>R</sup>, Leu | This study |
| pGADT7-AD-hTRAILR2<sub>DD</sub> | Human TRAILR2 death domain in pGADT7-AD, Amp<sup>R</sup>, Leu | This study |
| pGEX-4T-1 | Low copy number N-terminal glutathione-S-transferase fusion vector, Amp<sup>R</sup> | GE Healthcare |
| pGEX-SseK3 | *sseK3* from *S. Typhimurium* SL1344 in pGEX-4T-1, Amp<sup>R</sup> | This study |

**Table 3.**
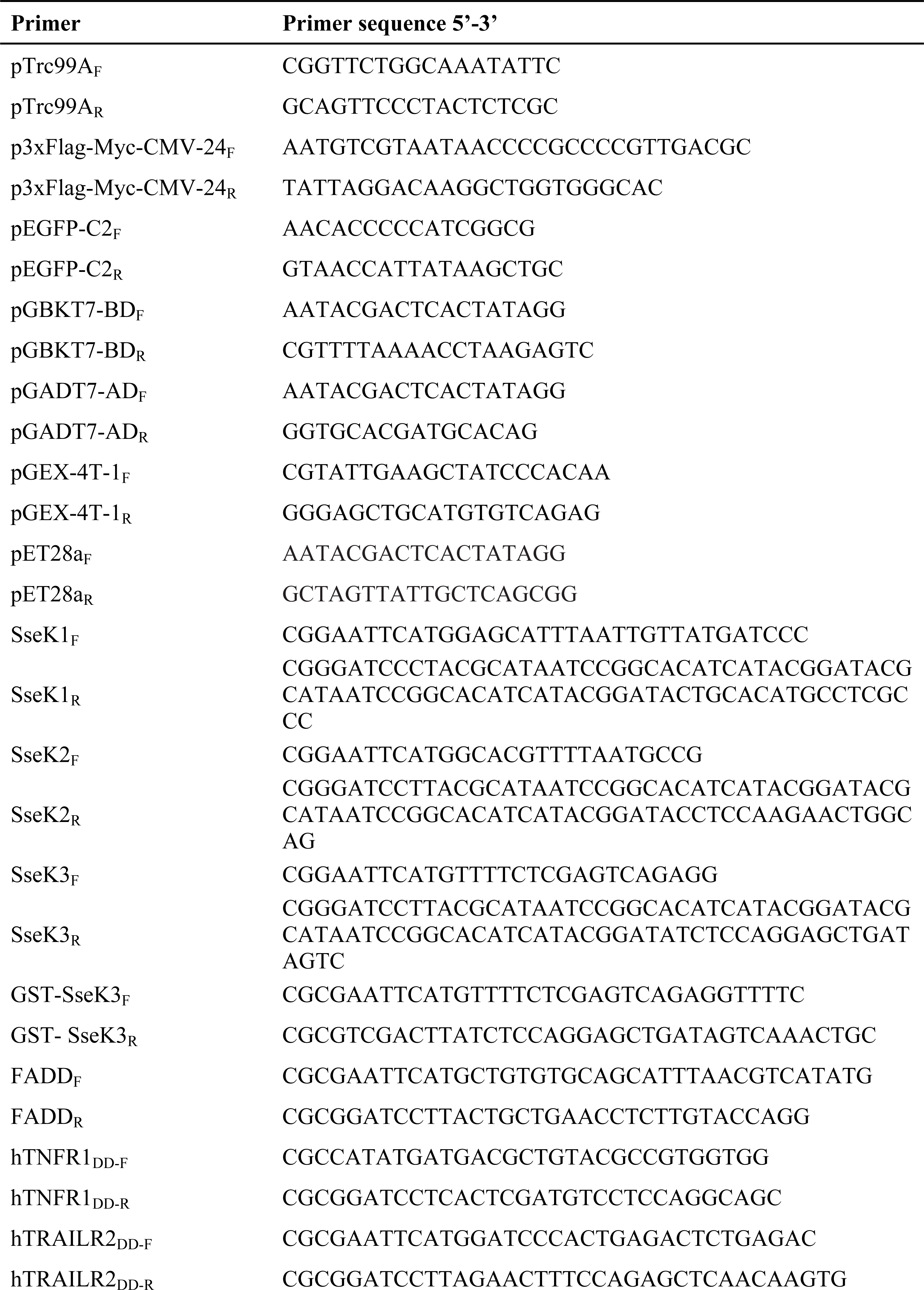
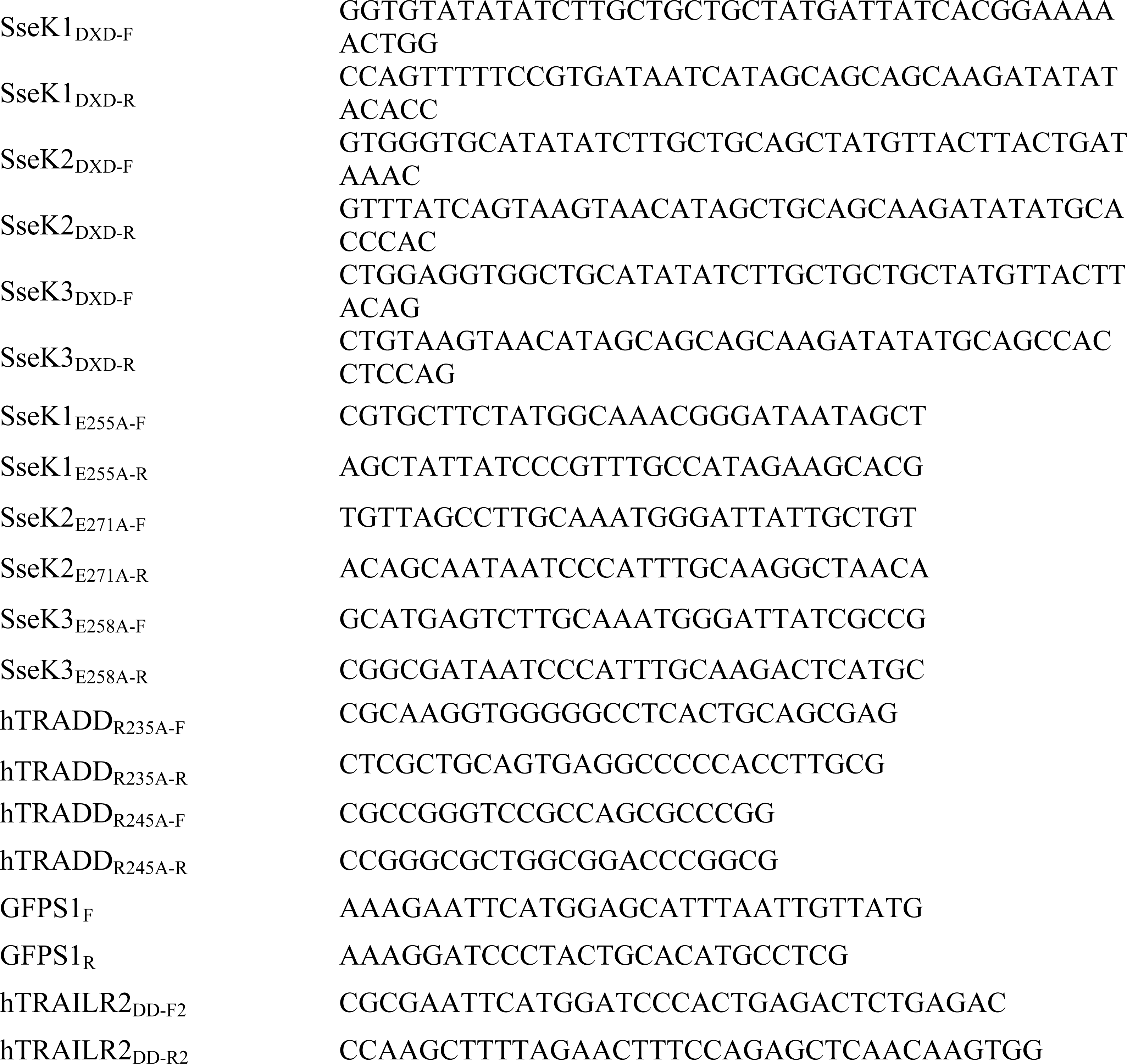
Primers used in this study.

### Overexpression of SseK1 alters the sites of glycosylation within the death domain of TRADD

A previous report demonstrated that TRADD carrying the amino acid substitution Arg^233^ was Arg-GlcNAcylated at comparable levels to wild-type TRADD when co-expressed ectopically with GFP-SseK1 in mammalian cells [9]. Our data indicated that mTRADD was glycosylated at Arg^243^ when SseK1 was overexpressed, but that Arg^233^ was the site of glycosylation when SseK1 was expressed at native levels during infection. The equivalent sites in human TRADD (hTRADD) are Arg^235^ and Arg^245^. To confirm these findings, we generated single and double Arg^235^ and Arg^245^ mutants of hTRADD and expressed these ectopically in HEK239T cells co-transfected with pEFGP-SseK1. Both Flag-hTRADD_R235A_ and Flag-hTRADD_R245A_ were Arg-GlcNAcylated at levels comparable to native Flag-hTRADD, while modification of the double mutant Flag-hTRADD_R235A/R245A_ was significantly reduced (Fig. 3A). To identify further possible sites of modification, Flag-hTRADD was enriched from cells co-transfected with pEGFP-SseK1 by anti-Flag immunoprecipitation, subjected to tryptic digestion and analysed by LC-MS (Fig. 3B, Supplementary Table 4). Under these conditions, we detected several further sites of modification at Arg^239^, Arg^278^, and Arg^224^, suggesting that over-expression broadened the range of possible glycosylation sites within the single substrate. The anti-Flag enrichment and LC-MS approach was repeated with Flag-hTRADD_R235A,_ Flag-hTRADD_R245A,_ and Flag-hTRADD_R235A/R245A_ variants (Supplementary Table 4). We detected different patterns of Arg-GlcNAcylation for each of these mutant Flag-TRADD variants, suggesting that deletion of a preferred glycosylation site caused a shift in site specificity despite comparable protein levels, as determined by MS analysis (Supplementary Table 5, Supplementary Fig 3). These data indicated that SseK1 was capable of modifying a range of arginine residues when expressed ectopically, and that over expression of SseK1 and related effectors may not replicate natural effector activity.

**Figure 3.**
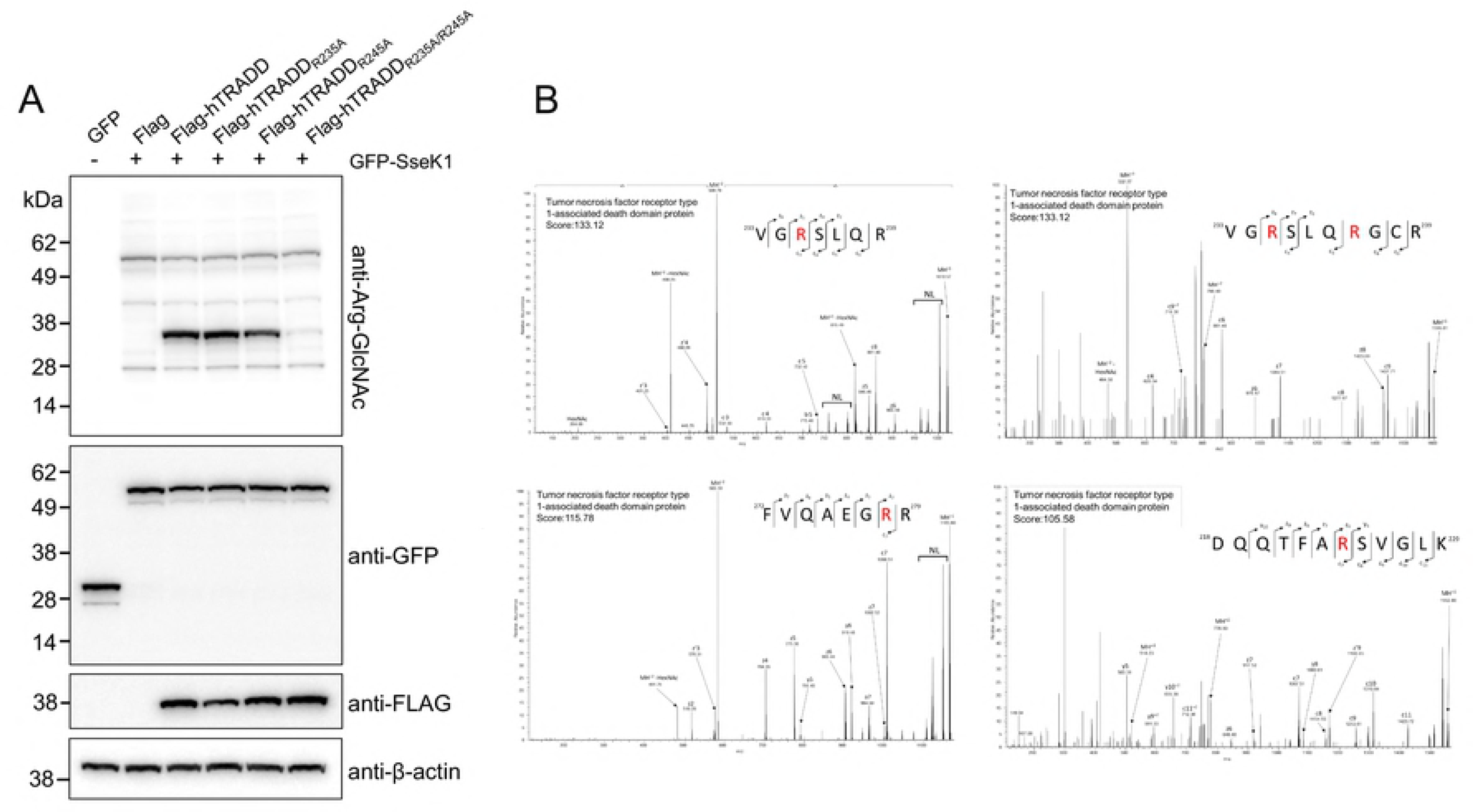
**Mutagenesis of putative SseK1 glycosylation sites of TRADD.** (A) Immunoblot showing Arg-GlcNAcylation of ectopically expressed Flag-hTRADD or Flag-hTRADD mutants in HEK293T cells co-transfected with pEGFP-SseK1. Cells were harvested for immunoblotting and detected with anti-ArgGlcNAc, anti-GFP, and anti-Flag antibodies. Antibodies to β-actin were used as a loading control. Representative immunoblot of at least three independent experiments. (B) Manually curated EThcD spectra of Arg-GlcNAcylated Flag-hTRADD enriched by anti-Flag immunoprecipitation following ectopic expression in HEK293T cells and co-transfection with pEGFP-SseK1. Various observed sites of Arg-GlcNAcylation are highlighted in red, and presented alongside corresponding M/Z values and observed Andromeda scores. Within MS/MS spectra NL denote neutral loss associated ions.

**Table 4.**
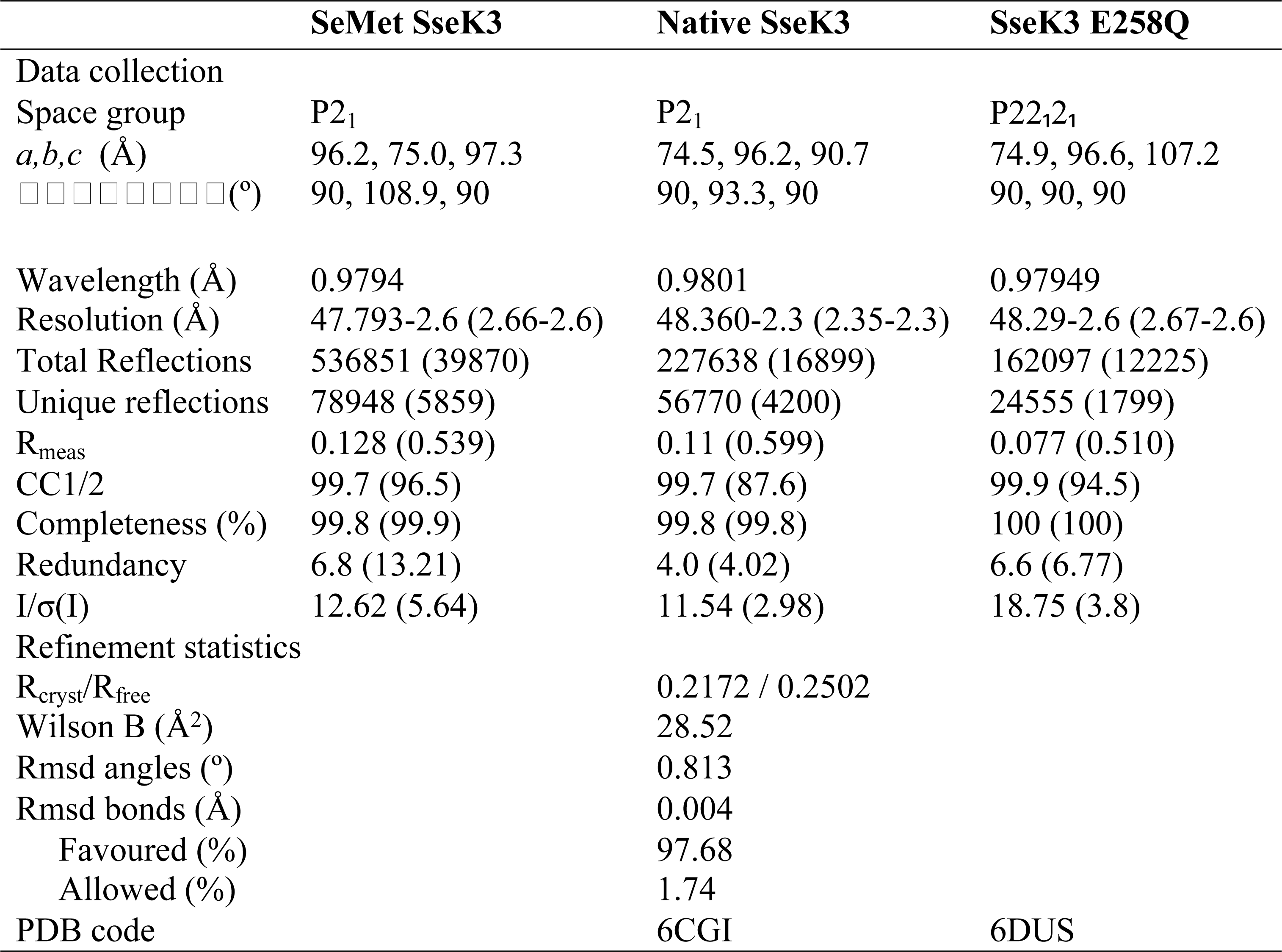
Data collection and structure refinement statistics.

### SseK3 glycosylates a conserved arginine residue in the mammalian death receptors TNFR1 and TRAILR

Similar to SseK1, we found that overexpression of SseK3 greatly increased arginine glycosylation activity relative to the levels generated by the bacterium during wild type *S.* Typhimurium infection (Fig. 1B). Hence, we applied our peptide enrichment strategy to identify Arg-GlcNAcylated substrates in the presence of native levels of SseK3. Arg-GlcNAcylated peptides were enriched from RAW264.7 cells infected with either a double *ΔsseK12* mutant or a triple *ΔsseK123* mutant, and we applied label free MS based quantification to screen for glycosylation events in a non-biased manner (Fig. 4A, Supplementary Table 6). Under these conditions, SseK3 modified specific arginine residues of mouse TNFR1 (mTNFR1) and TRAILR (mTRAILR), both death domain-containing receptors of the TNF superfamily [15]. Arg-GlcNAcylation was observed in all three biological replicates of cells infected with the *ΔsseK12* deletion mutant, while no Arg-GlcNAcylation was detected in cells infected with the *ΔsseK123* deletion mutant (Fig. 4B, Supplementary Table 6, Supplementary Fig. 4). Glycosylation of mTRAILR occurred at Arg^293^ while mTNFR1 was glycosylated at Arg^376^. An alignment of protein sequences demonstrated that these sites correspond to a conserved arginine in the death domains of both proteins (Figs. 4C and D). No other Arg-GlcNAcylated peptides were detected under these conditions, suggesting that TRAILR and TNFR1 were the preferred substrates of SseK3.

**Figure 4.**
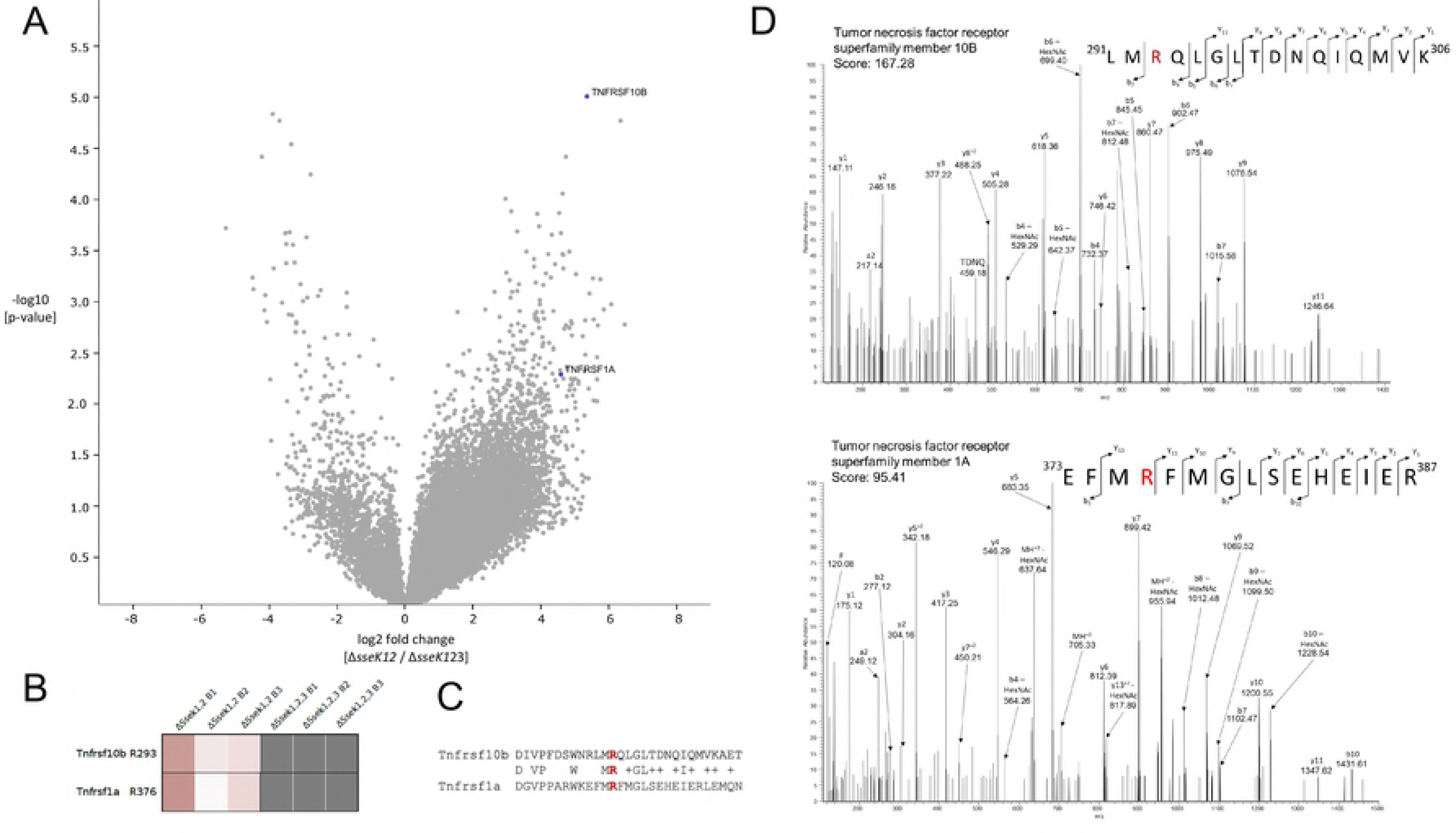
**Identifying substrates of SseK3 by Arg-GlcNAc peptide enrichment.** (A) Label-free quantification of Arg-GlcNAc peptide immunoprecipitated from RAW264.7 cells infected with *S.* Typhimurium Δ*sseK123* or Δ*sseK12*. Arginine-glycosylated peptides are presented as a volcano plot depicting mean log2 ion intensity peptide ratios of Δ*sseK123* versus Δ*sseK12* plotted against logarithmic *t* test *p* values from biological triplicate experiments. Arg-GlcNAcylated peptides are annotated by gene name and shaded blue. (B) Heat map showing observed ion intensity of Arg-GlcNAcylated peptides between biological triplicates. (C) Partial sequence alignment showing observed GlcNAcylated arginine residue is conserved between identified substrates. (D). Manually curated HCD spectra of arginine glycosylated TNFRSF10B/TNFR1 (upper) and TNFRSF1A/TRAILR (lower). Observed sites of Arg-GlcNAcylation are highlighted in red, and presented alongside corresponding M/Z values and observed Andromeda scores.

To determine if SseK3 also modified human TRAILR, the Flag tagged death domain of hTRAILR2 (Flag-hTRAILR2_DD)_ was enriched by anti-Flag immunoprecipitation from HEK293T cells cotransfected with pEGFP-SseK3. Of the four human isoforms of TRAILR, we focussed on hTRAILR2 as it shows the strongest sequence similarity to mTRAILR [16]. Using the anti-Arg-GlcNAc antibody, we detected Arg-GlcNAcylation of Flag-hTRAILR2_DD_ by GFP-SseK3 but not GFP-SseK3_E258A_ (Fig. 5A). Similarly, Flag-hTNFR1_DD_ was Arg-GlcNAcylated by GFP-SseK3 but not GFP-SseK3_E258A_ (Fig. 6A).

**Figure 5.**
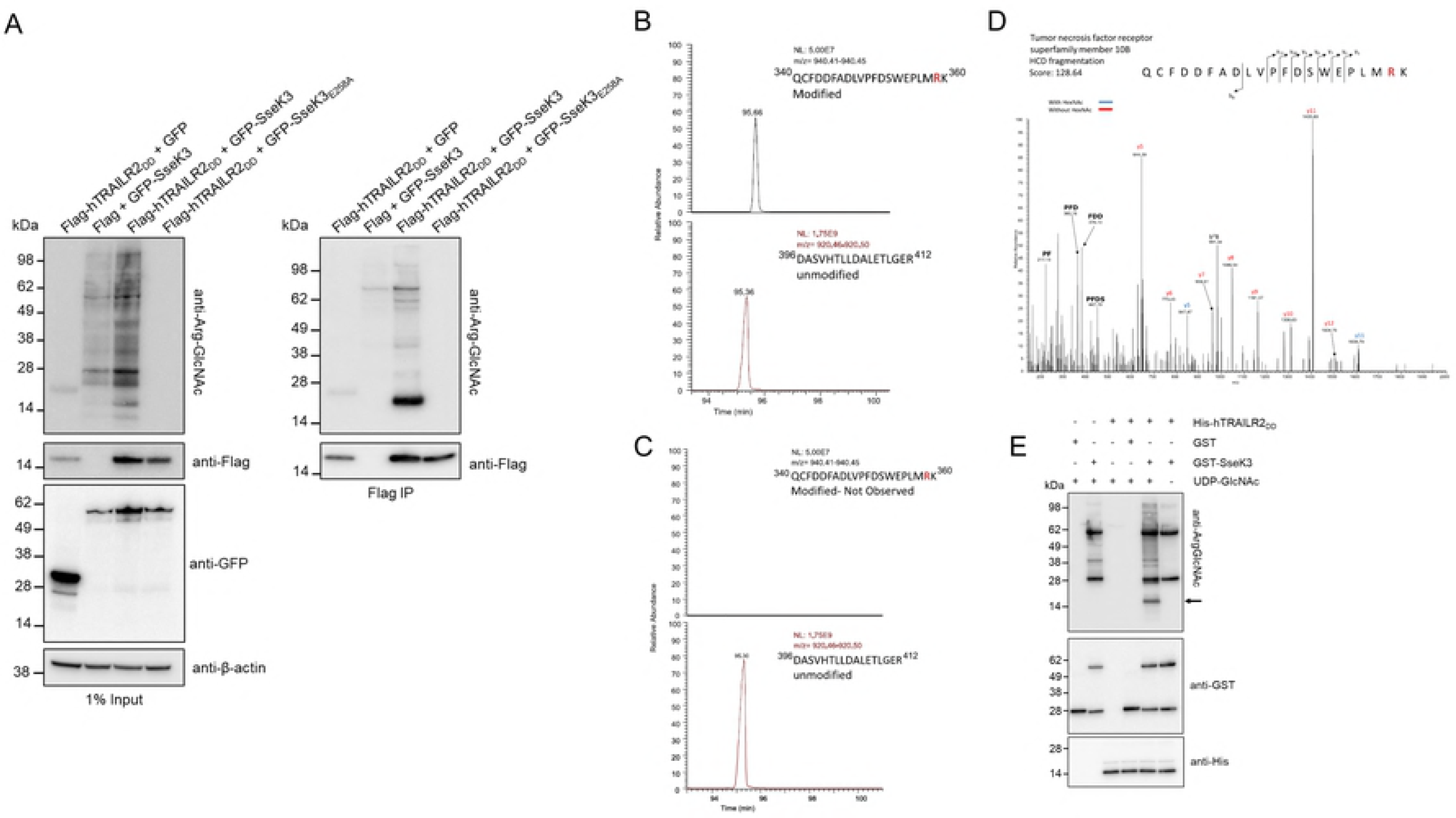
***In vitro*** **validation of host substrate modifications by SseK3.** (A) Immunoblot of inputs and immunoprecipitates (IP) of anti-Flag immunoprecipitations performed on lysates of HEK293T cells co-transfected with pFlag-hTRAILR2_DD_ and pEGFP-SseK3 or pEGFP-SseK3_E258A_. Proteins were detected with anti-Arg-GlcNAc, anti-GFP and anti-Flag as indicated. Antibodies to β-actin were used as a loading control. Representative immunoblot of at least three independent experiments. (B) LC-MS analysis of tryptic digest derived from co-incubation of recombinant His-hTRAILR2_DD_ and GST-SseK3 in the presence of UDP-GlcNAc. (C) LC-MS analysis of tryptic digest fractions derived from co-incubation of recombinant His-hTRAILR2_DD_ and GST-SseK3 with no sugar donor. (D) HCD fragmentation of recombinant His-hTRAILR2_DD_ incubated with GST-SseK3 and UDP-GlcNAc, Andromeda score 167.28. (E) Immunoblot of recombinant His-hTRAILR2_DD_ and GST-SseK3 following co-incubation at 37°C for 5 hours. Proteins were detected with anti-Arg-GlcNAc, anti-GST, and anti-His antibodies as indicated. Arrow indicates Arg-GlcNAcylated His-hTRAILR2_DD_. Representative immunoblot of at least three independent experiments.

**Figure 6.**
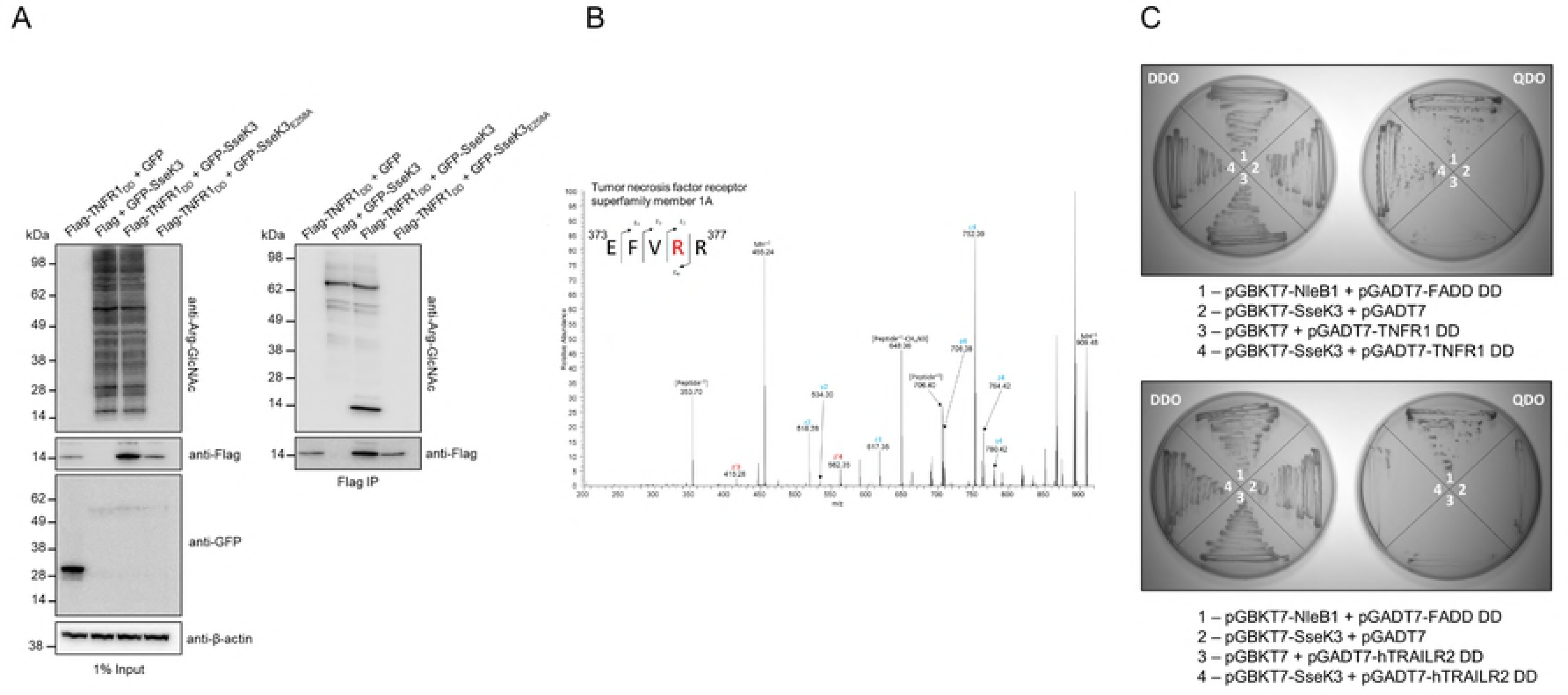
***In vitro*** **binding studies of SseK3 and Arg-GlcNAcylation of human TNFR1**. (A) Immunoblot of input and immunoprecipitate (IP) of anti-Flag immunoprecipitations performed on lysates of HEK293T cells co-transfected with pFlag-hTNFR1_DD_ and pEGFP-SseK3 or pEGFP-SseK3_E258A._ Proteins were detected with anti-Arg-GlcNAc, anti-GFP and anti-Flag as indicated. β-actin detection was used as a loading control. Representative immunoblot of at least three independent experiments. (B) EThcD fragmentation of Flag-hTNFR1_DD_ enriched from HEK293T cells by anti-Flag immunoprecipitation following co-transfection with pEGFP-SseK3, peptide confirmed by manual annotation. (C) *S. cerevisiae* Y2HGold co-transformed with pGBKT7-SseK3 and pGADT7-hTRAILR2 DD or pGADT7-hTNFR1 DD and plated onto selective media to select for plasmid carriage (DDO) or to select for protein-protein interactions (QDO). *S. cerevisiae* Y2HGold co-transformed with pGBKT7-NleB1 and pGADT7-FADD DD was used as a positive control for protein-protein interactions. Self-activation by the bait or prey fusion proteins was discounted by co-transformation of *S. cerevisiae* Y2HGold with pGADT7 and pGBKT7-SseK3 or co-transformation with pGBKT7 and pGADT7-hTNFR1 DD or pGADT7-hTRAILR2 DD.

To validate the specific site of modification, recombinant GST-SseK3 was incubated with HishTRAILR2_DD_ in the presence of the sugar donor UDP-GlcNAc, subjected to tryptic digestion and analysed by LC-MS (Supplementary Table 7 and 8). We detected hTRAILR2_DD_ modified at Arg^359^ (Fig. 5B, 5C), equivalent to the modification of mTRAILR at Arg^293^ described above. In the absence of UDP-GlcNAc, the modification of hTRAILR2_DD_ was not observed (Fig 5D). For both experiments the unmodified peptide ^396^DASVHTLLDALETGER^412^ derived from mTRAILR was monitored as an internal control and showed comparable ion intensity within the sample input which was consistent with the total observed protein levels between samples (Supplementary Table 8). Within these digests, we also observed evidence of self Arg-GlcNAcylation of SseK3, even without the addition of UPD-GlcNAc (Supplementary Table 7, Supplementary Fig 5 to 8). Immunoblots using anti-Arg-GlcNAc antibodies confirmed the glycosylation of hTRAILR2_DD_ in the presence of GST-SseK3 and UDP-GlcNAc as well as the apparent self-modification of SseK3 (Fig. 5E).

*In vitro* validation of hTNFR1 Arg-GlcNAcylation by GST-SseK3 was complicated by the insoluble nature of recombinant His-hTNFR1_DD_. Instead, Flag-hTNFR1_DD_ was enriched from HEK293T cells by anti-Flag immunoprecipitation following co-transfection with pEGFP-SseK3 (Fig. 6A). Protein samples were subjected to tryptic digestion ahead of analysis by LC-MS and Arg^376^ was confirmed as the preferred site of GlcNAcylation of Flag-hTNFR1_DD_ (Fig. 6B, Supplementary Table 9)

To explore the strength of the protein-protein interactions between SseK3 and the novel substrates, TNFR1 and TRAILR, the yeast two-hybrid system was used to detect protein-protein interactions. Auxotrophic yeast strains were co-transformed to express SseK3 and the death domain of hTNFR1 (hTNFR1_DD_). Yeast expressing SseK3 and hTNFR1_DD_ grew when plated on selective media, indicating a stable interaction between SseK3 and the hTNFR1_DD_ (Fig 6C). In contrast, yeast co-transformed to express SseK3 and the death domain of hTRAILR2 (hTRAILR2_DD_) did not grow on selective media, suggesting the interaction between these two proteins is either weaker, transient or that the death domain alone was insufficient for binding (Fig 6C).

Further to understanding the biology of SseK3, we observed that HA tagged SseK3 showed distinct Golgi localisation following *Salmonella* infection, as reported previously for ectopic expression [9]. Interestingly, this localisation did not require the glycosyltransferase activity of SseK3, suggesting that SseK3 may contain a novel Golgi targeting motif (Fig. 7).

**Figure 7.**
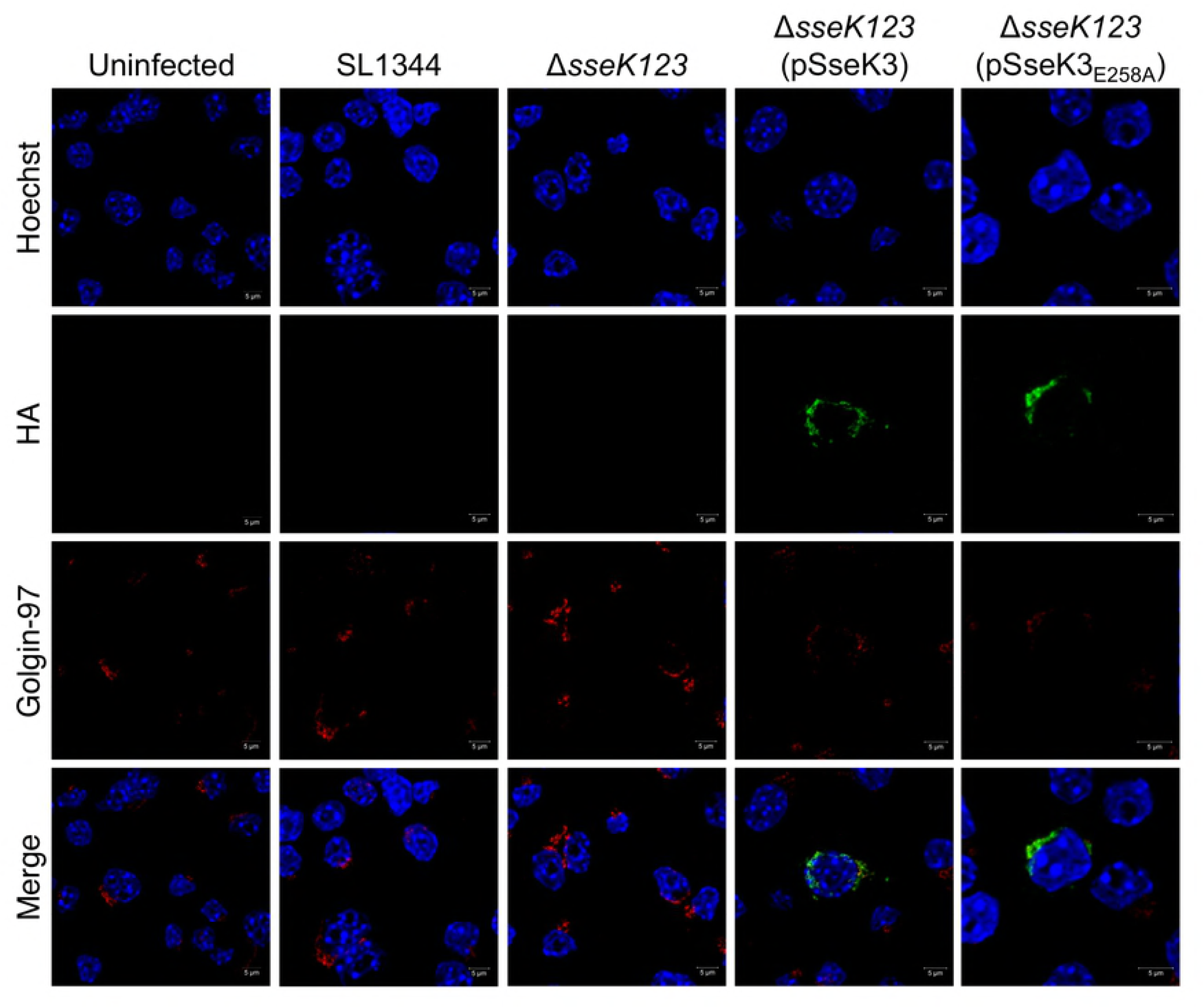
**Golgi localisation of SseK3.** Representative immunofluorescence fields of RAW264.7 cells infected with derivatives of *S.* Typhimurium SL1344 as indicated. The Δ*sseK123* triple mutant was complemented with a plasmid encoding HA-tagged SseK3 or catalytically-inactive HA-tagged SseK3_E258A_. Overexpression of effectors was induced during host cell infection by the addition of 1 mM IPTG. Cells were fixed and permeabilized 20 h after infection. The Golgi apparatus was visualised using anti-Golgin-97 antibodies (red). HA-tagged effector proteins were visualised using anti-HA antibodies (green). Cell nuclei were visualised with Hoechst stain (blue).

### Crystal structure of SseK3

To get a better understanding of the catalytic mechanism of SseK3 and related Arg-GlcNAc glycosyltransferases and the importance of the E258 residue, we solved the structure of SseK3(25-335) co-crystallized with UDP-GlcNAc and the E258Q mutant with UDP-GlcNAc. The enzyme belongs to the α/β class and has a core GT-A glycosyltransferase fold, with a central six-stranded β-sheet (Fig. 8A). There are two α-helices parallel to strands β1-β3 and two other helices parallel to the other three β-strands but on the opposite side of the β-sheet. An α-helical hairpin is inserted within the long connection between strands β3 and β4 of the central β-sheet that is somewhat separated from the rest of the protein and distant from the active site (Fig. 8A, B). Although the enzyme was co-crystallized with UDP-GlcNAc, only UDP is visible in the electron density of the native protein and there is no metal ion present (not added in the crystallization). UDP is located within a large, L-shaped groove with uridine filling the end of the short arm and the diphosphate extending to the middle corner of the L. The C-terminus of SseK3 is near the groove but the last five residues are disordered. In the SseK3 (E258Q) mutant, which crystallized in a different space group and in the presence of Mg^2+^ and UDP-GlcNAc, the C-terminus is ordered and extends over the UDP binding site, partially covering it from the solvent. The SseK3 E258Q mutant retained some glycosyltransferase activity since UDP-GlcNAc was also hydrolysed and only UDP was clearly visible in the crystal. However, an additional density was present nearby at the bottom of the long arm of L-groove and we have modelled GlcNAc into this density. In our structures, the UDP uracil O2 forms a hydrogen bond with the backbone amide of Phe53 while the uracil N3 atom forms a hydrogen bond with the backbone carbonyl oxygen of the same Phe53 (Fig. 8C). As well, the sidechains of Arg55 and Arg59 are close to the uracil O4. Moreover, the uracil ring forms a π-stacking interaction with Trp52 and Phe190. The ribose also forms hydrogen bonds to the protein, namely, C2 hydroxyl to the carbonyl of Gln51 and to the hydroxyl of Tyr224, while the C3 hydroxyl is hydrogen bonded to the backbone amide of Ala227. In the presence of Mg^2+^ ion (in E258Q mutant), both phosphates ligand the ion together with Asp228 (from the DxD^228^ motif), Asp325 and Ser327 (Fig. 8D). In addition, the α-phosphate hydrogen bonds to OG of Ser333 and β-phosphate hydrogen bonds to OG of Ser333 and NE1 of Trp334. In the absence of the metal ion (in the native structure) UDP still binds to the enzyme and uridine makes the same contacts. The phosphates are shifted ~2 Å away from the DxD motif and make no close contacts with the protein. In this structure, Tyr334 is disordered.

**Figure 8.**
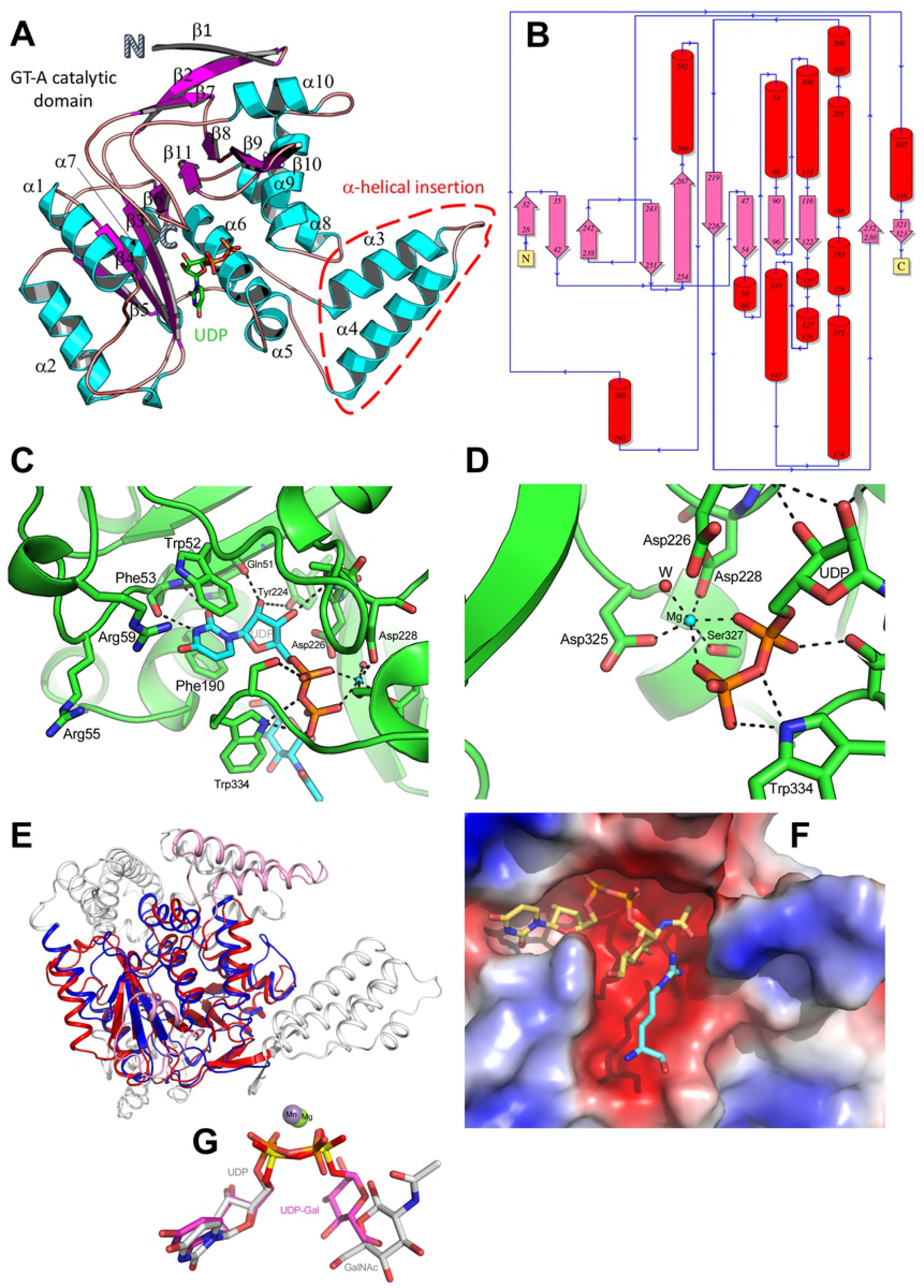
**SseK3 structure.** (A) Cartoon representation of the SseK3·UDP complex. The structure of SseK3 is shown with the bound UDP drawn as sticks. SseK3 displays a GT-A fold and contains an -helical insertion (marked by a dashed line). (B) Topology diagram of SseK3 with helices shown as red cylinders and strands as pink arrows. (C) Coordination of UDP in the E258Q active site. Residues in SseK3 are shown in green and the UDP moiety is shown in cyan. Hydrogen bonds are shown as black dashed lines. Arg59 is directed toward the uracil while Arg55 is mobile and assumes different conformations in various structures. (D) Coordination of the Mg^2+^ ion and the hydrogen bonds to the diphosphate in the E258Q active site. The Mg^2+^ ion is shown as a cyan sphere. (E) Superposition of SseK3 (red) and GT44 family *Clostridium difficile* toxin A (TcdA) glucosyltransferase domain (PDB code 3SRZ, blue). Toxin A domain is larger than SseK3 and the segments that have no correspondence in SseK3 are painted gray. Two segments of SseK3 without correspondence in toxin A are painted pink. (F) The UDP-GlcNAc and arginine are modelled into the active site of SseK3. SseK3 is represents as a solvent accessible surface coloured by the electrostatic potential (red – negative, blue – positive). UDP_GlcNAc was taken from structure 3SRZ and placed in SseK3 based on the position of UDP in the SseK3 crystal structure. Arginine was positioned in the long arm of the groove and can be easily accommodated. In this position, the arginine could accept GlcNAc from UDP-GlcNAc. The UDP moiety occupies the short arm of the L-shaped groove and an acceptor arginine is modelled into the long arm. (G) The superposition of UDP and GlcNAc from SseK3(E258Q) (white carbons) with UDP-Glc from toxin A glucosyltransferase domain (pink carbons).

Comparison of all independent molecules from crystal structures we determined showed that they superimpose with root-mean-squares deviation (rmsd) of 0.4-0.65 Å. The segments that deviated the most were the tip of the α-helical insertion mini-domain (aa 143-158) and the C-terminus, which is either well-ordered covering the bound UDP or completely disordered.

## Discussion

Elucidating the biochemical activity of bacterial effector proteins, their preferred host substrates, and contribution to virulence, remains a major priority of host-pathogen research to help elucidate basic processes that are manipulated by highly evolved pathogens, and to provide for new opportunities to disrupt disease causing processes. A number of studies have described putative substrates of the *Salmonella* Typhimurium effectors SseK1 and SseK3 [8, 9, 11, 12] but these studies have relied substantially on in vitro experiments using recombinant proteins or overexpression of the effectors in mammalian cells. Here, we identified the substrates of SseK1 and SseK3 when expressed at native levels during *S.* Typhimurium infection of RAW264.7 cells.

Although previous reports had suggested both TRADD and FADD were substrates of SseK1 [8, 9, 11], our data suggested that TRADD is the preferred substrate of SseK1, as we could not detect modification of FADD during *S.* Typhimurium infection. TRADD plays a key role in the activation of canonical NF-κB signaling leading to pro-inflammatory cytokine secretion (reviewed in [17, 18]) and programmed cell death via TNF induced apoptosis or necroptosis [19]. Previous work has suggested that SseK1 plays a role in inhibiting both NF-κB activation and necroptotic cell death in infected macrophages [9], and so it is likely that *S.* Typhimurium employs SseK1 to inhibit TNF signaling as required. Despite this, single deletion mutants of *S.* Typhimurium have not clearly demonstrated a requirement for SseK1 in mouse infection models [5, 20, 21]. We speculate that SseK1 acts in concert with other effectors *in vivo* to achieve significant inhibition of NF-κB signaling during *S.* Typhimurium infection.

Although SseK3 had been reported to interact with TRADD and TRIM32 [9, 12], here we found that the preferred substrates of SseK3 were TNFR1 and TRAILR, both members of the mammalian TNF receptor superfamily. TNFR1 responds to stimulation by extracellular TNF, and initiates a signalling cascade culminating in either inflammatory cytokine production or programmed cell death, as described above. Similarly, extracellular TRAIL binds the membrane-associated receptors TRAILR1 [22], TRAILR2 [23], TRAILR3 [24], and TRAILR4 [25]. However, only TRAILR1 and TRAILR2 contain full length death domains that stimulate a range of signaling events, including inflammatory cytokine production, extrinsic apoptotic cell death via caspases-8 and −3, necroptosis or even the promotion of cell survival via anti-apoptotic functions mediated by TRAF2 ubiquitination of caspase-8 [26, 27] (reviewed in [16, 28, 29]). This diversity of signaling outcomes complicates the interrogation of bacterial manipulation of TRAIL signaling. However, TRAIL deficient mice showed no difference in susceptibility to infection with *Salmonella* [30], which we confirmed (data not shown) suggesting that TNFR1 may be a more important target for bacterial inhibition in mammalian hosts. Consistent with this, TNFR1 deficient mice are highly susceptible to *Salmonella* infection [31]. Interestingly, a number of studies have implicated *TRAIL* polymorphisms with increased susceptibility to *Salmonella* infection in chickens [32, 33], suggesting TRAIL may activate an important immune defence pathway to *Salmonella* in poultry that SseK3 activity may block.

In addition to the Arg-GlcNAcylation of host substrates, we also observed the apparent self-modification of SseK1 and SseK3 when overexpressed. This observation, in conjunction with increased levels of Arg-GlcNAcylation seen during overexpression, suggested that many substrates may be non-authentically glycosylated when the effectors are overexpressed. These findings are similar to a previous report that suggested overexpression of NleB1 also results in Arg-GlcNAcylation of a broader range of substrates [13]. We note in particular the numerous two-component response regulators that were Arg-GlcNAcylated during overexpression of SseK1 (Fig 2A, Supplementary Table 1), perhaps suggesting a mechanism for effector-mediated regulation of two-component signaling outcomes. Surprisingly, these proteins also appeared to be Arg-GlcNAcylated by SseK1 and SseK3 during growth in LB (Supplementary Figure 9, Supplementary Table 10). Since these enrichment approaches did not provide information on the stoichiometry of these Arg-GlcNAcylation events, we attempted to interrogate this in *Salmonella* lysates without enrichment. However, this approach failed to detect any Arg-GlcNAcylation events, even for highly abundant proteins such as TufB and GlpK which we had found were modified following enrichment (Supplementary Figure 9). Overall this suggested that these events occur at low stoichiometry, and therefore the biological relevance of such glycosylation events is yet to be established.

Unlike SseK1, SseK3 shows a distinct subcellular localisation to the Golgi. As death receptors such as TNFR1 can be internalised and traffic with trans-Golgi vesicles [34], the localisation of SseK3 is consistent with modification of the receptors during infection. Interestingly, active glycosyltransferase activity was not required for Golgi localisation suggesting that the amino acid sequence of SseK3 may contain a novel Golgi targeting motif.

The crystal structure of SseK3 indicated that the enzyme has a GT-A glycosyltransferase fold and the closest structural homologs are enzymes from Carbohydrate-Active enZYmes Database (CAZy, http://www.cazy.org/) family GT44 and GT88, both containing bacterial retaining glucosyltransferases [35] (Fig. 8E). The structure showed the location of the substrate binding site as well as position of the active site residue, Glu258. The substrate binding groove has an L-shape and displays a negative electrostatic potential as would be expected to receive a positively charged arginine sidechain. The short arm of the groove is occupied by the UDP-GlcNAc and the long arm is most likely the binding site for the arginine sidechain. The E258Q mutant displayed some residual activity that prevented capturing the intact UDP-GlcNAc in the binding site but showed that in the absence of the acceptor, the GlcNAc is at least partially retained in the groove. Residual activity for the Glu-to-Gln mutation of a catalytic nucleophile was also observed in other retaining glucosyltransferases, for example, E317Q mutant of α-1,3-galactosyltransferase [36]. The mobile C-terminus likely forms a gate that allows for an easy access of UDP-GlcNAc to access the site and helps to retain it there until the acceptor arginine binds nearby and the transfer reaction is completed. To location and conformation of UDP observed in or structure superimposes very well with UDP-Glc from the crystal structure of one of the closest structural homologs, GT44 family *Clostridium difficile* toxin A (TcdA) glucosyltransferase domain (PDB code 3SRZ, [37]) (fig. 8G. Moreover, the position of GlcNAc observed in the SseK3(E258Q) structure was in the vicinity of the Glc of UDP-Glc in toxin A. In this transferase, the UDP-Glc binding site is also covered by a loop that is in a different conformation in the absence of UDP donor. The UDP conformation in SseK3 is also the same as UDP-GlcNAc in the structure of rabbit N-acetylglucosaminyltransferase I (PDB code 1FOA) [38]. In order to model a possible placement of the substrate arginine we have superimposed UDP-GlcNAc from the latter structure on UDP. The GlcNAc fits snugly into the groove and leaves space for the arginine, which we placed in the long arm of the substrate binding groove (Fig. 8E). It appears that this position of arginine is plausible for the transferase reaction. The role of the inserted mini-domain is unclear at the present time. Based on sequence alignment, this segment is present not only in SseK3 but also in SseK1/2 and NleB1/2 and we surmise that it plays a role in protein substrate recognition.

While this work was in progress, the structure of SseK3 crystallized under somewhat different conditions and in a different space group was published [39]. Comparison of these structures and the recently deposited structure of NleB2 (PDB code 5H5Y) shows flexibility of the ~10 C-terminal residues, particularly in the absence of UDP. When ordered, this segment covers the active and substrate binding sites, strengthening the hypothesis that these residues participate in UDP-GlcNAc and substrate binding and release.

In summary, here we identified the endogenous host substrates of SseK1 and SseK3 during *Salmonella* infection. The dominant substrate of SseK1 was the signaling adaptor TRADD, while SseK3 Arg-GlcNAcylated the death domains of both TRAILR and TNFR1 at a conserved arginine residue. This suggests that *Salmonella* utilises the SseK effectors to antagonise multiple components of death receptor pathways, thereby subverting inflammatory and cell death responses *in vivo*.

## MATERIALS AND METHODS

### Strains and growth conditions

The strains used in this study are listed in Table 1. Bacteria were grown with shaking at 37°C in Luria-Bertani (LB) broth in the presence of ampicillin (100 μg/ml), streptomycin (50 μg/ml), or kanamycin (100 μg/ml) when required.

### DNA cloning and purification

The plasmids and primers used in this study are listed in Tables 2 and 3, respectively. DNA-modifying enzymes were used in accordance with the manufacturer’s instructions (New England BioLabs). Plasmids were extracted using the QIAGEN QIAprep Spin Miniprep Kit. PCR products and restriction digests were performed using the Wizard SV Gel and PCR Clean-Up System (Promega). pTrc99A-SseK1, pTrc99A-SseK2, and pTrc99A-SseK3 were constructed by amplifying *sseK1, sseK2*, and *sseK3* from pEGFP-C2-SseK1, pEGFP-C2-SseK2, and pEGFP-C2-SseK3 using the primer pairs SseK1_F/R_, SseK2_F/R_, and SseK3_F/R_, respectively. The PCR product was digested with *Eco*RI and *Bam*HI and ligated into pTrc99A to produce a C-terminal 2x hemagglutinin tag fusion to SseK1, SseK2, and SseK3. Constructs were transformed into XL-1B cells, and verified by colony PCR and sequencing using the primer pair pTrc99A_F/R._ pEGFP-C2-SseK2 and pEGFP-C2-SseK3 were constructed by amplifying *sseK2* and *sseK3* from *S*. Typhimurium SL1344 genomic DNA using the primer pairs GFPS2_F/R_ and GFPS3_F/R_ respectively and AmpliTaq Gold DNA polymerase. The resultant PCR products were purified and ligated into pGEM-T-Easy vector at an insert:vector molar ratio of 3:1. The ligation reactions were transformed into XL-1 Blue cells and plated onto LA plates containing ampicillin and X-gal. Plasmids were extracted and digested with *Eco*RI and *Sal*I to release the bacterial genes, which were gel purified and ligated into pre-digested pEGFP-C2. The ligation reactions were then transformed into XL-1 Blue cells and colony PCR was performed using primers pEGFP-C2_F/R_ to select positive clones. The correct insert was confirmed by sequencing using the same primer pair. The primer pair hTRAILR2DD-_F/R_ was used to amplify the region encoding the death domain of TRAILR2 from HeLa cDNA (Sigma Aldrich). The resulting amplicon was gel purified and digested with *Eco*RI and *Bam*HI and ligated into pre-digested pGADT7. The ligation reactions were transformed into XL-1 Blue cells and plated on LA containing ampicillin. The correct insert was verified by colony PCR and sequencing with the primer pair pGADT7-AD_F/R_. pFlag-hTRAILR2_DD_ was constructed by amplifying the death domain of human *TNFRSF10B* from pGADT7- hTRAILR2_DD_ using the primer pair hTRAILR2_DD-F_/ hTRAILR2_DD-R_. The PCR product was digested with *Eco*RI and *Bam*HI and ligated into p3xFlag-*Myc*-CMV-24 to produce an N-terminal 3xFlag fusion to hTRAILR2_DD_. pFlag-hTRAILR2_DD_ was transformed into XL-1B cells, and verified by colony PCR and sequencing using the primer pair p3xFlag-Myc-CMV-24_F/R._ pGEX-4T-1-SseK3 was constructed by amplifying *sseK3* from pEGFP-C2-SseK3 using the primer pair GST-SseK3_F/R_. The PCR product was digested with *Eco*RI and *Sal*I and ligated into pre-digested pGEX-4T-1 to produce an N-terminal GST fusion to SseK3. pGEX-4T-1-SseK3 was transformed into XL-1B cells, and verified by colony PCR and sequencing using the primer pair pGEX-4T-1_F/R_. The region encoding the death domain of TNFR1 was amplified using the primer pair FLAGTNFR1DD_F/R_ and pGADT7-TNFR1_DD_ as template. The resulting amplicon was gel purified and digested with *Bgl*II and *Sal*I and ligated into pre-digested p3xFLAG-*Myc*-CMV-24 before transforming the reactions into XL-1 Blue cells. Colony PCR and sequencing were performed with the primer pair p3xFlag-Myc-CMV-24_F/R_ to ensure the correct insert has been ligated. pGBKT7-NleB1 was constructed by digesting pGBT9-NleB1 [7], which carries *nleB1* flanked in between the restriction sites *Eco*RI and *Bam*HI, and ligating into pGBKT7 digested with *Eco*RI and *Bam*HI. pGBKT7-SseK3 was constructed by amplifying *sseK3* from pEGFP-C2-SseK3 using the primer pair SseK3_F/R_. The PCR product was ligated into the cloning vector pGEM-T-Easy and the ligation reaction was transformed into XL-1 Blue cells. Transformants were then selected on LA containing ampicillin and X-gal. Plasmids were extracted and digested with *Eco*RI and *Sal*I to release the bacterial gene, which was then ligated into pre-digested pGBKT7 to create pGBKT7-SseK3. This plasmid was sequenced using the primer pair pGBKT7_F/R_. pGADT7-FADD_DD_, pGADT7-TNFR1_DD_, and pGADT7-hTRAILR2_DD_ were constructed by amplifying the death domain regions of human FADD, TNFR1, and TRAILR2 from HeLa cDNA using the primer pairs FADD_DD-F/R_, TNFR1_DD-F/R_, and hTRAILR2_DD-F/R_, respectively. Amplified FADD_DD_ was digested with *Eco*RI and *Bam*HI, TNFR1_DD_ was digested with *Nde*I and *Bam*HI, and hTRAILR2_DD_ was digested with *Eco*RI and *Bam*HI. Digested PCR products were ligated into pre-digested pGADT7 and transformed into XL-1 Blue cells. Constructs were verified by colony PCR and sequencing using the primer pair pGADT7_F/R_. For structural investigations, the segment corresponding to residues 25-335 of *sseK3* from *Salmonella Typhimurium (strain SL1344)* (Uniprot: A0A0H3NMP8) was cloned into vector pRL652, a derivative of vector pGEX-4T-1 (GE Healthcare) adapted for ligation-independent cloning. The construct contained a TEV-cleavable GST tag at the N-terminus.

### Site-directed mutagenesis

Site-directed mutagenesis of plasmid constructs was performed using the QuikChange II Site-Directed Mutagenesis kit (Stratagene, California, USA), according to the manufacturer’s instructions. pTrc99A-SseK1_DxD(229-231)AAA_, pTrc99A-SseK2_DxD(239-241)AAA_, and pTrc99A-SseK3_DxD(226-228)AAA_ were generated using pTrc99A-SseK1, pTrc99A-SseK2, or pTrc99ASseK3 as template DNA and amplified by PCR using the primer pairs SseK1_DxD-F/R_, SseK2_DxD-F/R_, or SseK3_DxD-F/R_, respectively. pTrc99A-SseK1_E255A_, pTrc99A-SseK2_E271A_, and pTrc99A-SseK3_E258A_ were generated using pTrc99A-SseK1, pTrc99A-SseK2, or pTrc99A-SseK3 as template DNA and amplified by PCR using the primer pairs SseK1_E255A-F/R_, SseK2_E271A-F/R_, or SseK3_E258A-F/R_, respectively. pFlag-hTRADD_R235A_ and pFlag-hTRADD_R245A_ were generated using pFlag-hTRADD as template DNA and amplified by PCR using the primer pairs hTRADD_R235A-F/R_ and hTRADD_R245A-F/R_, respectively. pFlag-hTRADD_R235A/R245A_ was generated using pFlaghTRADD_R235A_ as template DNA and amplified by PCR using the primer pair hTRADD_R245A-F/R_. All PCR products were digested with *Dpn*I at 37°C overnight before subsequent transformation into XL1-B cells. Plasmids were extracted and sequenced using the primer pairs pTrc99A_F/R_, p3xFlag-Myc-CMV-24_F/R_, or pEFGP-C2_F/R_, as required.

### Mammalian cell culture

HEK293T cells (human embryonic kidney 293 cells expressing the SV40 large T-antigen, source: ATCC^®^ CRL-3216) and RAW264.7 cells (murine leukemic monocyte-macrophage cells source: Richard Strugnell, University of Melbourne) were maintained in DMEM, low glucose with GlutaMAX™ supplement and pyruvate (DMEM (1X) + GlutaMAX(TM)-I) (Gibco, Life Technologies, NY, USA). Tissue culture media was further supplemented with 10% (v/v) heat-inactivated fetal bovine serum (FBS) (Thermo Fisher Scientific). Cells were maintained in a 37 °C, 5% CO_2_ incubator, and passaged to a maximum of 30 times. HEK293T cells were split when cells reached 80 to 90% confluency with 1 ml 0.05% Trypsin-EDTA (1X) (Gibco, LifeTechnologies) per 75 cm^2^ of tissue culture, then resuspended with 10 volumes of DMEM supplemented with FBS. RAW264.7 cells were physically detached with a cell scraper and further diluted in fresh DMEM supplemented with FBS.

### Infection of RAW264.7 cells

RAW264.7 cells were seeded to 24 well plates at a concentration of 3 x 10^5^ cells per well one day before infection. 10 ml LB broths containing appropriate antibiotic were inoculated with *Salmonella* strains and incubated at 37 °C overnight with shaking at 180 rpm. On the day of infection, the OD_600_ readings of the overnight culture were read and used to estimate bacterial counts. Cells were then infected at a multiplicity of infection (MOI) of 10. 24 well plates were centrifuged at 1500 rpm for 5 minutes at room temperature to promote and synchronise infection. Infected cells were incubated at 37 °C, 5% CO_2_ for 30 minutes. Culture media was replaced with media containing 100 μg/ml gentamicin (Pharmacia, Washington, USA), and cells were incubated at 37 °C, 5% CO_2_ for a further 1 hour. Culture media was replaced with media containing 10 μg/ml gentamicin, and where necessary, 1 mM IPTG, and cells were incubated at 37 °C, 5% CO_2_ to the required time, post infection.

### Immunoblotting

Cells were lysed in cold 1 x KalB lysis buffer (50 mM Tris-HCl pH 7.4, 150 mM NaCl, 1 mM EDTA, 1% (vol/vol) Triton X-100) supplemented with 2 mM Na_3_VO_4_, 10 mM NaF, 1 mM PMSF, and 1 x EDTA-free Complete protease inhibitor cocktail (Roche). Cell lysate was incubated for at least 30 minutes on ice, then cell debris was pelleted at 13000 rpm at 4 °C for 12 minutes. The soluble protein fraction was mixed with 4 x Bolt^®^ LDS sample buffer (Life Technologies) and DTT (Astral Scientific) to a final concentration of 50 mM. Proteins were boiled at 80 to 90 °C for 10 minutes, then loaded to Bolt^®^ 4-12% Bis-Tris Plus gels (Life Technologies) alongside SeeBlue^®^ pre-stained protein ladder (Life Technologies). Proteins were separated by electrophoresis using an XCell SureLock™ Mini-Cell system (Life Technologies) with 1 x Bolt^®^ MES SDS or 1 x Bolt^®^ MOPS SDS running buffer (Life Technologies), according to the manufacturer’s instructions. Following electrophoresis, proteins were transferred onto nitrocellulose membranes using the iBlot2^®^ gel transfer device (Life Technologies) and iBlot2^®^ nitrocellulose transfer stacks (Life Technologies), according to the manufacturer’s instructions. Membranes were blocked in 5% (w/v) skim milk in TBS (20 mM Tris, 50 mM NaCl, pH 8.0) with 0.1% (v/v) Tween 20 at room temperature for at least 1 hour with shaking at 60 rpm. Membranes were rinsed and washed in TBS Tween, then probed one of the following primary antibodies as required at 4 °C overnight with shaking at 60 rpm: rabbit monoclonal anti-ArgGlcNAc (Abcam), mouse monoclonal anti-HA (BioLegend), mouse monoclonal anti-GFP (Roche), mouse monoclonal anti-Flag M2-HRP (Sigma), or mouse monoclonal anti-β-actin (Sigma). Membranes were again rinsed and washed in TBS Tween, then probed with anti-mouse or anti-rabbit IgG secondary antibodies conjugated to horseradish peroxidase (PerkinElmer) diluted in TBS with 5% BSA (Sigma) and 0.1% Tween (Sigma) at room temperature for one hour with shaking at 60 rpm. Membranes were rinsed and washed in TBS Tween at room temperature for at least 45 minutes with shaking at 60 rpm. Antibody binding was detected using chemiluminescent substrates for horseradish peroxidase (HRP) (ECL western blotting reagents (GE Healthcare) or ECL Prime western blotting reagent (Amersham, USA), according to the manufacturer’s instructions, and visualised using an MFChemiBis imaging station.

### Transfection of HEK293T cells

HEK293T cells were transfected using FuGENE^®^6 transfection reagent (Promega), according to the manufacturer’s instructions. Cells were transfected one day after seeding to achieve 80 to 90% confluency. Transfection reagent was mixed with the reduced serum medium Opti-MEM^®^I (1X) + GlutaMAX(TM)-I (Gibco, Life Technologies), and incubated at room temperature for 5 minutes. Plasmid DNA was added at a transfection reagent:DNA ratio of 3:1, and incubated at room temperature for 25 minutes. The reaction was added to previously seeded cells and incubated at 37 °C, 5% CO_2_ for 16 to 24 hours.

### Immunoprecipitation of Flag-tagged fusion proteins

At required time points post-transfection, cells were lysed and the insoluble fraction removed as described above. Immunoprecipitation of Flag-tagged proteins was performed using Anti-Flag^®^ M2 Magnetic Beads (Sigma-Aldrich), according to the manufacturer’s instructions. Beads were first washed twice with lysis buffer, then mixed with the remaining soluble protein fraction and incubated rotating at 4 °C overnight. Following this, beads were washed three times with lysis buffer. Bound protein was eluted by incubating beads in 60 μl of 150 μg/ml Flag peptide (Sigma-Aldrich), rotating at 4 °C for 30 minutes. Eluate was mixed with LDS and DTT, and boiled at 80 to 90 °C for 10 minutes. Input and eluate samples were electrophoresed, then visualised by immunoblot as above.

### Yeast two hybrid assay

Yeast strain *S. cerevisiae* Y2H Gold (Clontech, California, USA) was transformed or cotransformed with plasmid DNA using the established lithium acetate method [40]. Transformants were plated to selective media as required to select for successful single or double transformation. When validating interactions between two proteins, transformants were subsequently plated to highly selective media. Briefly, *S. cerevisiae* Y2H Gold was streaked to YPDA and incubated at 30 °C for 3 days. Healthy colonies were used to inoculate 10 ml YPDA broth at a starting OD_600_ of 0.2, and incubated at 30 °C with shaking at 200 rpm to an OD^600^ of 0.6-0.8. The yeast culture was centrifuged at 4000 rpm for 7 minutes, and the pellet was resuspended in sterile distilled water, and centrifuged again. The yeast culture was then resuspended in 100 mM lithium acetate, vortexed thoroughly, and centrifuged again. The lithium acetate supernatant was removed, and yeast were resuspended in 400 mM lithium acetate, vortexed thoroughly, and centrifuged again. The lithium acetate supernatant was removed, and yeast were resuspended in polyethylene glycol (PEG 3350, Sigma-Aldrich), 1 M lithium acetate, salmon sperm ssDNA at a final concentration of 2 mg/ml, and plasmid DNA as appropriate. This reaction was incubated at 30 °C for 30 minutes, then subjected to heat shock at 42 °C for 20 minutes. The reaction was briefly centrifuged, and the supernatant removed. The yeast pellet was resuspended in distilled water and plated to both SD/Trp-Leu and SD/Trp-Leu-Ade-His selective media, then incubated at 30 °C for 3 days.

### *In vitro* glycosylation assay

His-tagged and GST-tagged fusion proteins were purified from bacterial cultures using Novagen His-Bind^®^ purification kit or Novagen GST-Bind™ purification kit, respectively, according to the manufacturer’s instructions. Briefly, plasmids encoding either 6 x His-tagged or GST-tagged fusion proteins were transformed into BL21 C43 (DE3) *E. coli*. Overnight cultures grown in LB with appropriate antibiotics were used to inoculate a 200 mL LB subculture (1:100) which was grown for 3 hours at 37 °C with shaking at 180 rpm to an optical density of 0.6. Subcultures were induced with 1 mM isopropyl-β-D-thiogalactopyranoside (IPTG; AppliChem, Darmstadt, Germany), and grown for a further 3 hours. Cultures were then centrifuged at 10000 rpm at 4 °C for 15 minutes, then resuspended in the appropriate resuspension buffer. Resuspended bacteria were lysed using an EmulsiFlex-C3 High Pressure Homogenizer (Avestin), according to the manufacturer’s instructions. Lysates were centrifuged at 13000 rpm at room temperature for 30 minutes, and proteins were purified from the soluble fraction by either nickel- or glutathione-affinity chromatography, according to the manufacturer’s instructions.

Protein concentrations were determined using a bicinchoninic acid (BCA) kit (Thermo Fisher Scientific). Recombinant proteins (approximately 1 μg) were incubated alone or together, and in the presence of 1 mM UDP-GlcNAc (Sigma-Aldrich). Reactions were made to a total volume of 80 μl in TBS (50 mM Tris, 150 mM NaCl, pH 7.6) supplemented with 10 mM MgCl_2_ and 10 mM MnCl^2^. Reactions were incubated at 37 °C for 4-5 hours. To detect *in vitro* glycosylation, reactions were either electrophoresed and probed by Western blot as above, or processed for mass spectrometry analysis as below.

### Tryptic digest of gel-separated proteins

Affinity purified proteins were separated using SDS-PAGE, fixed and visualized with Coomassie G-250 according to protocol of Kang *et al.* [41]. Bands of interest were excised and destained in a 50:50 solution of 50 mM NH_4_HCO_3_ / 100% ethanol for 20 minutes at room temperature with shaking at 750 rpm. Destained samples were then washed with 100% ethanol, vacuum-dried for 20 minutes and rehydrated in 50 mM NH_4_HCO_3_ plus 10 mM DTT. Reduction was carried out for 60 minutes at 56 °C with shaking. The reducing buffer was then removed and the gel bands washed twice in 100% ethanol for 10 minutes to remove residual DTT. Reduced ethanol washed samples were sequentially alkylated with 55 mM Iodoacetamide in 50 mM NH_4_HCO_3_ in the dark for 45 minutes at room temperature. Alkylated samples were then washed with two rounds of 100% ethanol and vacuum-dried. Alkylated samples were then rehydrated with 12 ng/µl trypsin (Promega) in 40 mM NH_4_HCO_3_ at 4 °C for 1 hour. Excess trypsin was removed, gel pieces were covered in 40 mM NH_4_HCO_3_ and incubated overnight at 37 °C. Peptides were concentrated and desalted using C18 stage tips [42, 43] before analysis by LC-MS.

### Enrichment of arginine-glycosylated peptides from infected cell lysate

Infected cells were washed three times in ice-cold PBS and lysed by scraping with ice-cold guanidinium chloride lysis buffer (6 M GdmCl, 100 mM Tris pH 8.5, 10 mM TCEP, 40 mM 2-Chloroacetamide) on a bed of ice according to the protocol of Humphrey *et al.* [44]. Lysates were collected and boiled at 95 ˚C for 10 minutes with shaking at 2000 rpm to shear DNA and inactivate protease activity. Lysates were then cooled for 10 minutes on ice then boiled again at 95 ˚C for 10 minutes with shaking at 2000 rpm. Lysates were cooled and protein concentration determined using a BCA assay. 2 mg of protein from each sample was acetone precipitated by mixing 4 volumes of ice-cold acetone with one volume of sample. Samples were precipitated overnight at −20 ˚C and then spun down at 4000 G for 10 minutes at 4 ˚C. The precipitated protein pellets were resuspended with 80% ice-cold acetone and precipitated for an additional 4 hours at −20 ˚C. Samples were spun down at 17000 G for 10 minutes at 4 ˚C to collect precipitated protein, the supernatant was discarded and excess acetone driven off at 65 ˚C for 5 minutes.

Dried protein pellets were resuspended in 6 M urea, 2 M thiourea, 40 mM NH_4_HCO_3_ and reduced / alkylated prior to digestion with Lys-C (1/200 w/w) then trypsin (1/50 w/w) overnight as previously described [45]. Digested samples were acidified to a final concentration of 0.5% formic acid and desalted with 50 mg tC18 SEP-PAK (Waters corporation, Milford, USA) according to the manufacturer’s instructions. Briefly, tC18 SEP-PAKs were conditioned with buffer B (80% ACN, 0.1% formic acid), washed with 10 volumes of Buffer A* (0.1% TFA, 2% ACN), sample loaded, column washed with 10 volumes of Buffer A* and bound peptides eluted with buffer B then dried. Peptide affinity purification was accomplished according to the protocol of Udeshi *et al.* [46], modified to allow for Arg-GlcNAc enrichment. Briefly, aliquots of 100 μl of Protein A/G plus Agarose beads (Santa Cruz, Santa Cruz CA) were washed three times with 1 ml of immunoprecipitation buffer (IAP, 10 mM Na_3_PO_4_, 50 mm NaCl, 50 mM MOPS, pH 7.2) and tumbled overnight with 10 μg of anti-Arg-GlcNAc antibody (ab195033, Abcam) at 4 ˚C. Coupled anti-Arg-GlcNAc beads were then washed three times with 1 ml of 100 mM sodium borate (pH 9) to remove non-bound proteins and cross-linked for 30 minutes rotating using 20 mM Dimethyl Pimelimidate (Thermo Fisher Scientific) in 100 mM HEPES, pH 8.0. Cross-linking was quenched by washing beads with 200 mM ethanolamine, pH 8.0, three times then rotating the beads in an additional 1 ml 200 mM ethanolamine, pH 8.0 for 2 hours at 4 ˚C. Beads were washed three times with IAP buffer and used immediately.

Purified peptides were resuspended in 1 ml IAP buffer and the pH checked to ensure compatibility with affinity conditions. Peptide lysates were then added to the prepared cross-linked anti-Arg-GlcNAc antibody beads and rotated for 3 hours at 4 °C. Upon completion antibody beads were spun down at 3000 G for 2 minutes at 4 °C and the unbound peptide lysates collected. Antibody beads were then washed six times with 1 ml of ice-cold IAP buffer and Arg-GlcNAc peptides eluted using two rounds of acid elution. For each elution round, 100 μl of 0.2% TFA was added and antibody beads allowed to stand at room temperature with gentle shaking every minute for 10 minutes. Peptide supernatants were collected and desalted using C18 stage tips [42, 43] before analysis by LC-MS.

### Identification of arginine-glycosylated affinity enriched peptides and Flag-tagged proteins using reversed phase LC-MS

Purified peptides prepared were re-suspend in Buffer A* and separated using a two-column chromatography set up composed of a PepMap100 C18 20 mm x 75 μm trap and a PepMap C18 500 mm x 75 μm analytical column (Thermo Fisher Scientific). Samples were concentrated onto the trap column at 5 μL/min for 5 minutes and infused into an Orbitrap Fusion™ Lumos™ Tribrid™ Mass Spectrometer (Thermo Fisher Scientific) at 300 nl/minute via the analytical column using a Dionex Ultimate 3000 UPLC (Thermo Fisher Scientific). 125 minutes gradients were run altering the buffer composition from 1% buffer B to 28% B over 90 minutes, then from 28% B to 40% B over 10 minutes, then from 40% B to 100% B over 2 minutes, the composition was held at 100% B for 3 minutes, and then dropped to 3% B over 5 minutes and held at 3% B for another 15 minutes. The Lumos™ Mass Spectrometer was operated in a data-dependent mode automatically switching between the acquisition of a single Orbitrap MS scan (120,000 resolution) every 3 seconds and Orbitrap EThcD for each selected precursor (maximum fill time 100 ms, AGC 5*104 with a resolution of 30000 for Orbitrap MS-MS scans). For parallel reaction monitoring (PRM) experiments the known tryptic Arg-modified sites of TRADD [8] and FADD [7] (Uniprot accession: B2RRZ7 and Q3U0V2 respectively) were monitored using the predicted m/z for the +2 and +3 charge states. Data-independent acquisition was performed by switching between the acquisition of a single Orbitrap MS scan (120000 resolution, m/z 300-1500) every 3 seconds and Orbitrap EThcD for each PRM precursor (maximum fill time 100 ms, AGC 5*104 with a resolution of 60000 for Orbitrap MS-MS scans).

### Mass spectrometry data analysis

Identification of proteins and Arg-glycosylated peptides was accomplished using MaxQuant (v1.5.3.1) [47]. Searches were performed against the Mouse (Uniprot proteome id UP000000589 – Mus musculus, downloaded 18-05-2016, 50306 entries), *Salmonella* Typhimurium SL1344 (Uniprot proteome id UP000008962- *Salmonella* Typhimurium SL1344, downloaded 18-05-2016, 4,657 entries) or human (Uniprot proteome id UP000005640-*Homo sapiens*, downloaded 24/10/2013, 84,843 entries) proteomes depending on the samples with carbamidomethylation of cysteine set as a fixed modification. Searches were performed with trypsin cleavage specificity allowing 2 miscleavage events and the variable modifications of oxidation of methionine, N-Acetylhexosamine addition to arginine (Arg-GlcNAc) and acetylation of protein N-termini. The precursor mass tolerance was set to 20 parts-per-million (ppm) for the first search and 10 ppm for the main search, with a maximum false discovery rate (FDR) of 1.0% set for protein and peptide identifications. To enhance the identification of peptides between samples the Match Between Runs option was enabled with a precursor match window set to 2 minutes and an alignment window of 10 minutes. For label-free quantitation, the MaxLFQ option within Maxquant [48] was enabled in addition to the re-quantification module. The resulting protein group output was processed within the Perseus (v1.4.0.6) [49] analysis environment to remove reverse matches and common protein contaminates prior. For LFQ comparisons missing values were imputed using Perseus and Pearson correlations visualized using Matlab R2015a (http://www.mathworks.com).

### Immunofluorescence microscopy

RAW264.7 cells were seeded onto coverslips at a density of 10^5^ cells per well 24 hours prior to infection. Cells were infected with *S.* Typhimurium SL1344 or indicated mutant strains for 18 hours. Infected cells were fixed in 4% (w/v) paraformaldehyde (Sigma) in PBS for 12 minutes on ice, then washed 3 times with PBS, and incubated in NH_4_Cl 1:100 in PBS at room temperature for 20 minutes. Cells were washed twice with PBS, then incubated in 0.2% (v/v) Triton X-100 (Sigma) in PBS at room temperature for 3 minutes. Cells were washed three times with PBS, then blocked in 3% (w/v) BSA in PBS at room temperature for 30 minutes. Coverslips were incubated in the following primary antibodies for 1 hour at room temperature: rabbit monoclonal anti-HA (Cell Signaling) and mouse monocloncal anti-Golgin-97 (Invitrogen), used 1:200 in PBS with 3% BSA. Coverslips were then incubated in the following secondary antibodies for 30 minute at room temperature: anti-rabbit AlexaFluor 488 (Invitrogen) and anti-mouse AlexaFluor 568 (Invitrogen), used in 1:2000 in PBS with 3% BSA. Coverslips were subsequently incubated in Hoechst staining solution (Sigma) for 10 minutes, diluted 1:5000 in PBS. Coverslips were mounted onto microscope slides using Prolong Gold mounting medium (Life Technologies) and images were acquired using a Zeiss confocal laser scanning microscope with a 100x EC Epiplan-Apochromat oil immersion objective.

### Protein expression and purification for structural studies

The pRL652 plasmid containing GST-TEV-Ssek#(25-335) was transformed into BL21(DE3) strain. For protein expression a 15 mL overnight culture in LB was inoculated into one liter of terrific broth media supplemented with 100 µg/ml of ampicillin. The inoculated cultures were grown at 37^°^C until the OD_600_ reached 1.0. The cultures were transferred to 18^°^C, induced with 1mM of isopropyl β-D-1-thiogalactopyranoside (IPTG), left overnight to grow and harvested by centrifugation at 9,110 x g for 7 minutes.

The cell pellet was resuspended in lysis buffer (50 mM Tris-HCl buffer pH 8.0, 10% glycerol, and 0.1% Triton X) and the cells were lysed in a cell disruptor (Constant Systems Ltd., Northants, United Kingdom). Cell debris were removed by centrifugation at 28,965 x g for 30 minutes. The supernatant was loaded on 10 mL Glutathione-Superflow resin (Clontech) column equilibrated with standard buffer (20 mM Tris pH 8.0 and 150 mM NaCl). The column was washed with 5 column volumes of standard buffer. The beads were then incubated with TEV protease (33 μg/mL) in 30 mL of standard buffer for overnight at room temperature. The flow-through containing cleaved SseK3 was collected, concentrated to 30 mg/mL with the Millipore centrifugal filter with a molecular weight cut-off of 10,000 for crystallization trials. The E258Q mutant was generated using KOD Hot Start DNA Polymerase (Sigma-Aldrich) with mutagenic primers and WT plasmid as a template according to the manufacturer’s instructions. The E258Q mutant was purified using the same protocol.

The seleno-methionine (SeMet) derivative of SseK3 was expressed in auxotrophic *E. coli* strain B834(DE3). A 50 mL overnight culture in medium A (M9 medium, trace elements, glucose, MgSO_4_, CaCl_2_, Biotin, 50 mg/mL methionine and thiamin) was used to inoculate 1 L of medium A. All media used were supplemented with 100 µg/ml of Ampicillin. The cells were grown with shaking at 37^°^C until OD_600_ reached 1.0. The cells were pelleted at 4^°^C and then resuspended in 1 L of media A without methionine. The culture was further incubated for 4 hours at 37^°^C and 50 mg of SeMet was then added. After 30 minutes of incubation, the cultures were then induced with 1 mM of IPTG and continued to grow for additional 10 h at 18^°^C. The cells were harvested by centrifugation at 9,100 x g for 7 min. The SeMet-labelled protein was purified the same way as the native protein.

### Protein Crystallization

Initial crystals were obtained by screening using commercial and in-house screens in a 96-well plate format. The crystallization was setup using Gryphon crystallization robot (Art Robbins Instruments, Sunnyvale, CA). The best crystals were obtained using hanging-drop vapor diffusion method at 20^°^C. 1 μL of protein solution supplemented with 3 mM of UDP-GlcNAc was mixed with 1 μL of reservoir solution containing 0.5 M NaCl, 0.1 M Tris pH 8.5, 18% PEG 3350 and 5% MPD and suspended over 0.5 mL of reservoir. These crystals displayed P2_1_ space group symmetry. The SeMet-containing SseK3 crystallized at slightly different conditions and these crystals had the same space group symmetry as the native crystals but they differed in cell dimensions (Table 4). The best crystals of the SseK3(E258Q) mutant in comlplex with UDPGlcNAC and Mg^2+^ were obtained at somewhat different conditions, 0.5 M NaCl, 0.1 M Tris pH 8.5, 12% PEG 3350 and 5% MPD supplemented with 6 mM UDP-GlcNAc and 6 mM MgCl_2_. They displayed P2_1_2_1_2_1_ space group symmetry (Table 4).

### Data collection and structure determination

For data collection, the crystals were soaked briefly in a cryo-protecting solution containing 30% MPD and 70% reservoir solution and flash-cooled in liquid nitrogen. Diffraction data were collected at the Canadian Light Source (CLS) beam line 08B1-1 for the native crystal and 08ID-1 for the SeMe-labelled crystal. Data were processed using XDS program [50] with the AutoProcess script [51]. The structure was solved by single-wavelength anomalous dispersion (SAD) method using program AutoSol in Phenix program suite [52]. The initial model was refined using Phenix software interspaced with manual rebuilding using COOT [53]. The initial model of the SseK3 molecule was placed in the context of the native dataset by molecular replacement. There are four molecules in the asymmetric unit. The refinement continued with the Phenix software until convergence was reached with R_work_=0.2172 and R_free_=0.2502. The final model contains residues 27-330 for molecule A, 28-329 for molecule B, 26-329 for molecule C and D. Each molecule contains bound UDP. There are 292 solvent molecules. The structure of the E258Q mutant was solved by molecular replacement and contains two molecules in the asymmetric unit. The refinement converged with R_work_=0.177 and R_free_=0.225. The final model contains residues 26-335 in molecule A and 28-335 in molecule B, UDP, Mg^2+^ and GlcNAc bound to each molecule, three molecules of TRIS, five molecules of methyl-pentanediol and 141 water molecules. Data collection and refinement statistics are shown in Table 4. The coordinates and structure factors were deposited in the PDB data bank with accession numbers 6CGI and 6DUS.

## Acknowledgments

The authors are indebted to Jürg Tschopp for the gift of pFlag-TRADD. Research described in this paper was performed using beamline 08B1-1 and 08ID-1 at the Canadian Light Source, which is supported by the Canada Foundation for Innovation, Natural Sciences and Engineering Research Council of Canada, the University of Saskatchewan, the Government of Saskatchewan, Western Economic Diversification Canada, the National Research Council Canada, and the Canadian Institutes of Health Research.

## Supporting Information Legends

**Supplementary Table 1. Arginine-GlcNAcylated peptide pull downs from SseK1 overexpression during infection.** A total of 133 unique arginine-GlcNAcylated peptides corresponding to 113 localized Arg-GlcNAc sites observed from triplicate biological infections with Δ*sseK123* over-expressing either SseK1 or inactivated SseK1_E255A_. The site of arginine-GlcNAcylation is provided with 113 sites localized, as defined by a localization score of >0.75, and an additional 35 sites unable to be confidently localized (localization score of <0.75). For assigned arginine glycopeptides and arginine glycosylation sites, the protein, gene, score, ion intensity, and biological replicate in which the identity was observed are provided.

**Supplementary Table 2. Arginine-GlcNAcylated peptides pull downs from SSeK1 endogenous levels infections.** A total of 8 unique arginine-GlcNAcylated peptides corresponding to 11 localized Arg-GlcNAc sites observed from triplicate biological infections with Δ*sseK123* or Δ*sseK23*. The site of arginine-GlcNAcylation is provided with 11 sites localized, as defined by a localization score of >0.75, and an additional 35 sites unable to be confidently localized (localization score of <0.75). For assigned arginine glycopeptides and arginine glycosylation sites, the protein, gene, score, ion intensity, and biological replicate in which the identity was observed are provided.

**Supplementary Table 3: iBAQ based analysis of ΔsseK2/3 infections colon inputs for Arg-GlcNAc enrichment.** For the 5664 proteins observed from RAW264.7 infected with Δ*sseK23*, iBAQ intensity values generated using Maxquant are provided. For assigned proteins, the iBAQ values, score, summed ion intensity, number of MS/MS events, LFQ values and protein name gene, and biological replicate in which the identity was observed are provided.

**Supplementary Table 4. Arginine-GlcNAcylated peptides and sites from co-transfection with Flag-hTRADD variants and pEGFP-SseK1.** A total of 15 unique arginine-GlcNAcylated peptides are observed from hTRADD upon transfection of pEGFP-SseK1. Across the different Flag-hTRADD variants, different patterns of arginine-GlcNAcylation are observed with 13 sites of arginine-GlcNAcylation identified. For assigned arginine glycopeptides and sites, the protein, gene, score, ion intensity, and biological replicate in which the identity was observed are provided.

**Supplementary Table 5. Proteins identified from co-transfection with Flag-hTRADD variants and pEGFP-SseK1.** Protein analysis of Flag-hTRADD variants in co-transfected samples. For assigned proteins, the LFQ values, score, summed ion intensity, number of MS/MS events and protein name gene, and biological replicate in which the identity was observed are provided.

**Supplementary Table 6. Arginine-GlcNAcylated peptides pull downs from SSeK3 endogenous levels infections.** A total of 14 unique arginine-GlcNAcylated peptides corresponding to 6 localized Arg-GlcNAc sites observed from triplicate biological infections with Δ*sseK123* or Δ*sseK12*. The site of Arginine-GlcNAcylation provided with 6 sites localized, as defined by a localization score of >0.75, and 1 additional sites unable to be confidently localized (localization score of <0.75). For assigned arginine glycopeptides and arginine glycosylation sites the protein, gene, score, ion intensity, and biological replicate in which the identity was observed are provided.

**Supplementary Table 7. Arginine-GlcNAcylated peptides and sites from in vitro assays of hTRAIL and SseK3.** A total of 61 unique arginine-GlcNAcylated peptides corresponding to 15 Arg-GlcNAc sites within Ssek3 and hTRAIL were observed within in-vitro Arg-GlcNAcylation assays. For assigned arginine glycopeptides and arginine glycosylation sites, the protein, gene, score, ion intensity, and in vitro condition in which the identity was observed are provided.

**Supplementary Table 8. Proteins identified from** ***in vitro*** **assays of hTRAIL and SseK3.** Protein analysis of in vitro assay samples. For assigned proteins, the LFQ values, score, summed ion intensity, number of MS/MS events and protein name gene, and biological replicate in which the identity was observed are provided.

**Supplementary Table 8. Proteins identified from** ***in vitro*** **assays of hTRAIL and SseK3.** Protein analysis of in vitro assay samples. For assigned proteins, the LFQ values, score, summed ion intensity, number of MS/MS events and protein name gene, and biological replicate in which the identity was observed are provided.

**Supplementary Table 9. Proteins identified from co-transfection of hTNFR1_DD_ and SseK3.** Protein analysis of co-transfection assay samples of hTNFR1_DD_. For assigned proteins, the LFQ values, score, summed ion intensity, number of MS/MS events and protein name gene, and biological replicate in which the identity was observed are provided.

**Supplementary Table 10. Arginine-GlcNAcylated of Salmonella proteins.** A total of 40 unique arginine-GlcNAcylated peptides were observed from triplicate biological experiments of stationary phase Δ*sseK123,* Δ*sseK12,* Δ*sseK23* and wild type SL1344. For assigned arginine glycopeptides and arginine glycosylation sites the protein, gene, score and biological replicate in which the identity was observed are provided.

**Supplementary Figure 1. Pearson correlation of SseK1 related pulldowns.** To assess the reproducibility of arginine-GlcNAcylation, pull down heat maps of observed peptides are provided. A) Δ*sseK/23* over-expressing either SseK1 or inactivated SseK1_E255A_. The mean correlation between each replicate is 0.74922. B) Δ*sseK123* or *Salmonella* Δ*sseK23* infections. The mean correlation between each replicate is 0.70054.

**Supplementary Figure 2. Observed proteome of** *Salmonella* **Δ*****sseK2/3*** **infection inputs.** During infection of RAW264.7 cells with Δ*sseK23*, 5664 proteins were observed across the input proteomes. Relative abundance of proteins based on the log_10_ iBAQ intensity values generated using Maxquant are plotted based on ranked order. The position in the rank order of death domain containing proteins and TNFRSF proteins are shown. Within the input of all samples TRADD is higher in relative abundance then FADD.

**Supplementary Figure 3. FLAG-hTRADD levels observed in co-transfection studies.** The LFQ values for all co-transfection of FLAG-hTRADD variants are shown. Comparable levels of TRADD were observed within all samples.

**Supplementary Figure 4. Pearson correlation of SseK3 related pulldowns.** To assess the reproducibility of arginine-GlcNAcylation pull down, heat maps of observed peptides are provided for Δ*sseK12* or Δ*sseK123* infections. The mean correlation between each replicate is 0.737509.

**Supplementary Figure 5. The identification of Arg-GlcNAcylation sites within recombinant SseK3.** Manually curated EThcD spectra confirming the glycosylation of arginine-335 within the SseK3.

**Supplementary Figure 6. The identification of Arg-GlcNAcylation sites within recombinant SseK3.** Manually curated EThcD spectra showing glycosylation of arginine-305 within SseK3.

**Supplementary Figure 7. The identification of Arg-GlcNAcylation sites within recombinant SseK3.** Manually curated EThcD spectra showing glycosylation of arginine-153 within SseK3.

**Supplementary Figure 8. The identification of Arg-GlcNAcylation sites within recombinant SseK3.** Manually curated EThcD spectra showing glycosylation of arginine-184 within SseK3.

**Supplementary Figure 9. Arg-GlcNAcylation within stationary phase** *Salmonella* **grown in LB.** A) Ion Intensity heatmap of Arg-GlcNAcylation sites identified from biological triplicate pulldown from Δ*sseK123,* Δ*sseK12,* Δ*sseK23* and wild type SL1344 during stationary phase growth in LB. All arginine-GlcNAcylated peptides were dependent on SseK1 and SseK3 B) Pearson correlation of Arg-GlcNAc pulldowns within LB grown strains of *Salmonella*. To assess the reproducibility of arginine-GlcNAcylation pull down heat maps of observed peptides are provided.

